# The Connexin 43 Carboxyl Terminal Mimetic Peptide αCT1 Prompts Differentiation of a Collagen Scar Matrix Resembling Unwounded Skin

**DOI:** 10.1101/2020.07.07.191742

**Authors:** Jade Montgomery, William J. Richardson, J. Matthew Rhett, Francis Bustos, Katherine Degen, Gautam S. Ghatnekar, Christina L. Grek, Spencer Marsh, L. Jane Jourdan, Jeffrey W. Holmes, Robert G. Gourdie

## Abstract

Phase II clinical trials have reported that acute treatment of surgical skin wounds with the therapeutic peptide αCT1 improves cutaneous scar appearance by 47% 9-months post-surgery – though mode-of-action remains unknown. Scar matrix structure in biopsies 2 to 6 weeks post-wounding treated topically with αCT1 or control treatments from human subjects, Sprague-Dawley rats, and IAF hairless guinea pigs were compared. The sole effect on scar structure in humans was that αCT1-treated scars had less alignment of collagen fibers relative to control wounds, a state that resembles unwounded skin. This more random alignment was recapitulated in both animal models, together with transient increases in collagen density, although the guinea pig was found to more closely replicate the pattern of response to αCT1 in human scars, compared to rat. Fibroblasts treated with αCT1 *in vitro* showed decreased directionality and an agent-based computational model parameterized with fibroblast motility data predicted collagen alignments in simulated scars consistent with that observed experimentally in human and the animal models. In conclusion, αCT1 prompts decreased directionality of fibroblast movement and the generation of a 3D collagen matrix post-wounding that is similar to unwounded skin – changes that correlate with long-term improvement in scar appearance.

## Introduction

Cutaneous injury can result in scars that are not only unsightly, but also stiff, painful, and susceptible to further injury.^1^ Over 300 million surgical procedures are performed each year, often resulting in significant scarring.^2^ Together with scars caused by accidental or malicious injury there is a major unmet clinical need for safe and effective anti-scarring therapy. Unfortunately, to date there is no single standard-of-care exists for promoting wound resolution to minimize scar formation and encourage phenotypes associated with uninjured skin. Topically applied silicone patches have demonstrated modest efficacy in the treatment of scars, including hypertrophic and keloid scars.^3, 4^ However, the US Food and Drug Administration is yet to approve a treatment that reliably eliminates and/or inhibits excess deposition of fibrotic tissue that can accompany normal skin wound healing.^5–7^

Connexin 43 (Cx43) is a transmembrane channel protein that is expressed in both the epidermal and dermal layers of the skin, which has been shown to have key assignments in the dermal injury response.^8–16^ Regulated reduction of Cx43 expression in wounded skin is essential for normal wound healing,^17^ with Cx43 levels being dysregulated and overexpressed in chronic non-healing wounds.^18, 19^ Genetically reducing the overall level of Cx43 in normal mice resulted in accelerated wound re-epithelialization and wound closure, increased dermal fibroblast activity, and enhanced expression of extracellular matrix (ECM) remodeling factors.^20, 21^ Topical treatment of murine cutaneous wounds with a Cx43 antisense oligonucleotide has been reported to reduce inflammation and scar formation.^14^

Alpha Connexin Carboxy-Terminus 1 (αCT1) is a peptide mimic of the Carboxyl Terminus (CT) of Cx43, encompassing a class II PSD95/Dlg/ZO-1 (PDZ) binding motif that interacts with the PDZ 2 domain of Zonula Occludens-1 (ZO-1).^8, 15, 22, 23^ In addition to interaction with ZO-1 PDZ2, αCT1 has been shown to directly interact with the H2 domain of Cx43.^24^ Mouse cardiac injury models have indicated this interaction prompts phosphorylation of a serine at position 368 on Cx43 – a post-translational modification linked to reduced activity of Cx43-formed membrane channels.^13, 16, 25–28^ αCT1 interaction with ZO-1 may also reduce the density of Cx43 hemichannels in the membrane by prompting sequestration into gap junction plaques.^23^ At the macroscopic level, cutaneous woundhealing studies in mice and pigs have supported a role for αCT1 in increasing skin wound closure rate, tempering inflammatory neutrophil infiltration, reducing granulation tissue area during the first 10-20 days (sub-acute phase) of healing and improving the long-term mechanical properties of cutaneous scars.^8, 15, 29, 30^

Consistent with preclinical observations, a Phase II clinical trial determined that αCT1 improved scar visual appearance by 47% 9 months post-surgery relative to within-patient controls following laparoscopic surgery.^31^ This long-term improvement showed a trend of progressive emergence over the study period, with a modest 12.9 % augmentation over control observed at 3 months, but no apparent difference between αCT1 and control scars at 1 month. In the present study, we took advantage of an earlier vehicle-controlled Phase I clinical trial evaluating the safety and tolerability of αCT1 involving 49 healthy subjects in which scar tissue was biopsied from the wound site 29 days following cutaneous injury.^15^ These samples provided the opportunity to conduct histological evaluation of granulation tissue (scar progenitor tissue) in healed αCT1 -treated versus vehicle-treated control wounds within the same patient. Our analyses support a role for αCT1 in the organization of ECM in the early scar, wherein peptide-treated wounds showed a less aligned three-dimensional (3D) order of collagen bundles 4 weeks post-injury, a finding that was recapitulated in two animal models. By examining fibroblast behavior *in vitro*, and applying agent-based mathematical modeling, we provide evidence that the peptide’s effect on differentiating granulation tissue may be mediated by αCT1 promoting more random patterns of cell motility.

## Results

### Histological Characterization of Skin Samples from the Phase I Clinical Trial

Clinical testing of αCT1 was performed on 49 healthy volunteers in a randomized, double-blind Phase I study in Switzerland. On Day 1, a biopsy punch was used to create a circular 5 mm full-thickness wound of unblemished skin underneath both arms (Fig. 1A). Wounds were internally randomized within each patient and one wound was treated with αCT1 in a hydroxyethylcellulose gel, while the other was treated with a vehicle control gel, enabling within-patient comparisons. Participating individuals were randomized into four cohorts, wherein wounds were treated with vehicle gel or 20 μM (Cohort 1), 50 μM (Cohort 2), 100 μM (Cohort 3), or 200 μM (Cohort 4) of αCT1-containing gel. The gel was applied immediately after injury and again 24 hours later. On day 29 of the study, circular 2 mm full-thickness biopsies were collected from the healed granulation tissue formed at each of the two wound sites on each patient. The 100 μM dose of αCT1 applied to treated wounds in Cohort 3 is the therapeutic dose that was subsequently used in the Phase II clinical trial on 92 patients, wherein this dosage was associated with a clinically meaningful 47% improvement in scar appearance 9 months following treatment.^31^

**Figure 1.**
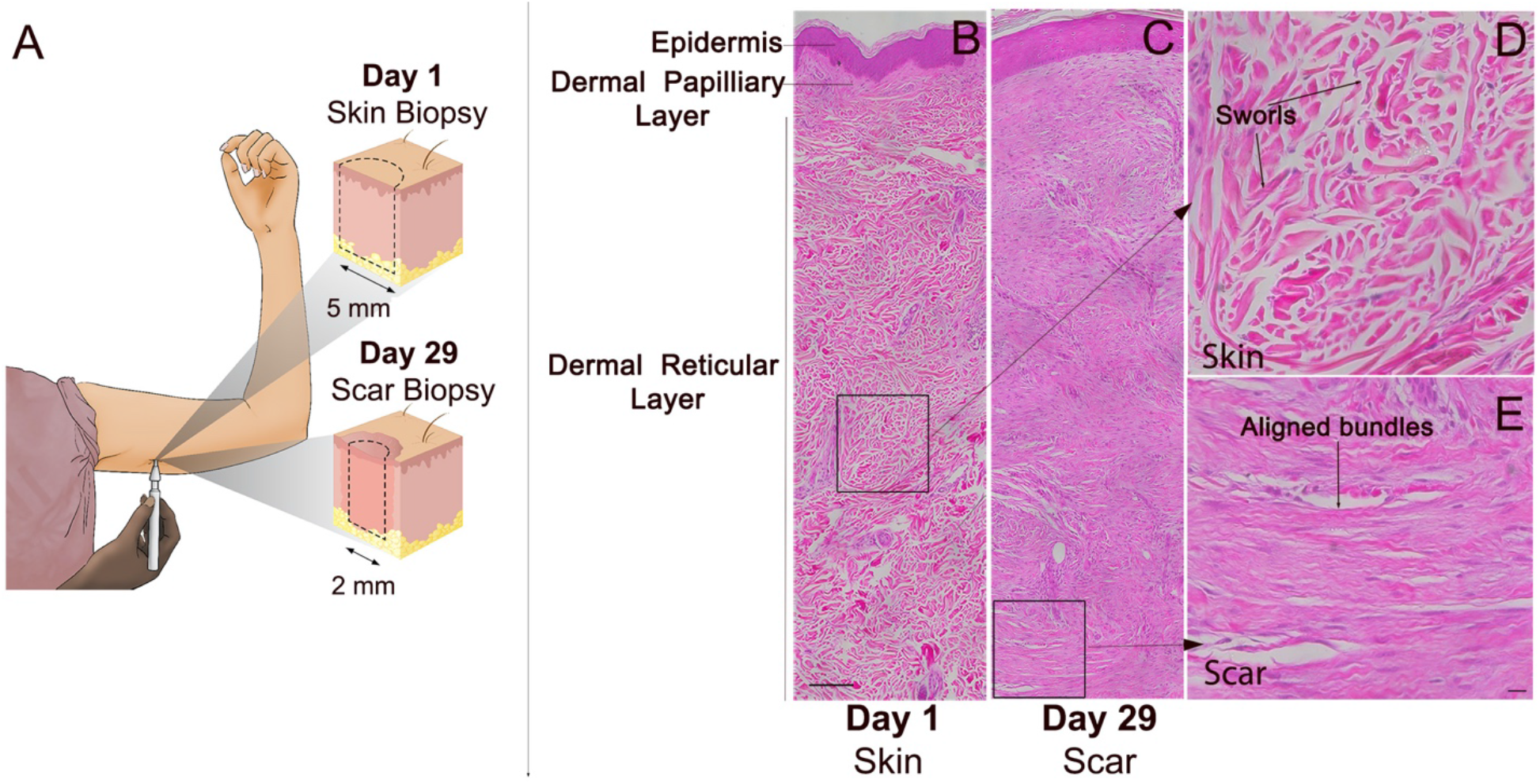
Phase I clinical trial on effects of αCT1 on cutaneous wounding in healthy humans. **A)** Phase I clinical trial skin sampling strategy. A biopsy punch was used to create a circular 5 mm full-width biopsy of unblemished skin underneath the upper arm. A circular 2 mm full-width biopsy of scar tissue was re-sampled at the same location 29 days later. H&E histochemical stained sections of: **B)** Unwounded skin sampled on day 1 from a vehicle control wound. The region boxed in the dermal layer in (B) is shown at higher magnification in (D) and **C)** Scar tissue re-sampled at the same location as that in (B) on day 29 of the study. The region boxed in the dermal layer of (C) is shown at higher magnification in panel E. Higher magnification images from boxed regions in (B) and (C) showing characteristic histoarchitecture of: **D)** Unwounded skin – including randomly organized dermal collagen bundles non-aligned with respect to epidermis and arranged in large sworls and **E)** Cutaneous scar – displaying dermal collagen bundles that are larger, more aligned and not arranged into sworls. Scale bar = 100 μm.

Field stitching microscopy (20x objective) was used to reconstruct montages of skin/scar hematoxylin and eosin (H&E) stained biopsy sections from the Phase I study. The scans included the full width (5-8 mm) of unwounded skin and scar biopsies from both arms of all patients. Exemplar images of unwounded skin (biopsy taken at study start) and scar tissue taken 29 days later from the healed biopsy site on the same patient after application of vehicle control gel are shown in figures 1B and C, respectively. Notable differences between unwounded skin (Fig. 1B) and post-wounding scar at one month (Fig. 1C), include the epidermal layer with rete pegs invaginating into the dermal layer of unwounded skin, which were absent in the epidermal layer of the healing scar. Also apparent are distinct bundles of collagen-rich extracellular matrix (ECM) in the dermal reticular layer of unwounded skin, which were woven into irregularly ordered whorls (Figs. 1B & D). Neither the ECM bundles nor the larger whorls displayed a consistent orientation with respect to the epidermis in unwounded skin – being organized in random orientations in all 45 patients (e.g., Supplemental Fig. 1). By contrast, ECM bundles in the scar biopsies had a more dense array of fibers arranged into thick, aligned tracts that ran mostly parallel to the epidermal layer, particularly near the dermal base (Figs. 1C & E).

To quantitatively compare histological features of scar tissue from αCT1- and vehicle-treated wounds, epidermal and dermal parameters from each of the 29-day full-thickness H&E-stained samples were evaluated by investigators blinded to patient, treatment and cohort (Fig. 2). The variables measured included: epidermal thickness, dermal-epidermal junction length, epidermal length, epidermal rete peg number, sub-epidermal melanocyte density, dermal nuclear density, dermal blood vessel density and intensity of dermal eosin staining (% eosin). Sebaceous glands, hair follicles, and eccrine glands were conspicuous in unwounded skin, but hardly present in scar biopsies, regardless of treatment – and thus were not analyzed. To obtain information on whether these histological parameters varied with skin depth, the dermis on the full-width sections was divided into 4 equally spaced quartiles, the upper quartile being closest to the epidermis and lowest incorporating the interval most proximal to the base of the skin adjacent to the hypodermis (Fig. 2A). The only epidermal or dermal variable that showed a significant (p<0.05) treatment or treatment by depth effect was the normalized percentage of intensity of eosin staining (% eosin) in the dermis - a metric approximating the density of ECM present in the tissue (Fig. 2B).

**Figure 2.**
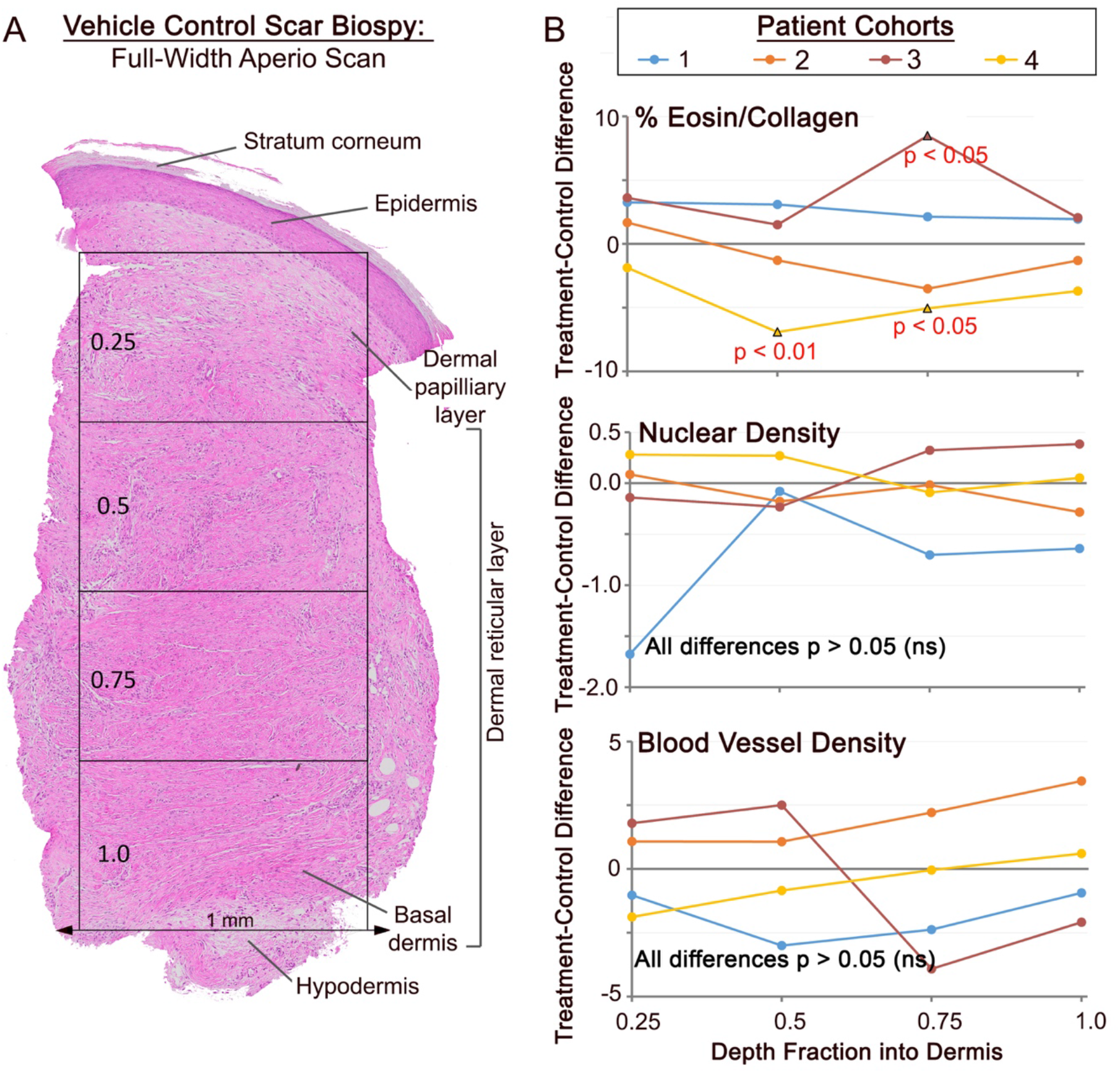
Initial histochemical study of αCT1 effects on human 29-day scar biopsies. **A)** Full-skin-width Aperio Imagescope (20x objective) montage of an H&E stained section of a day 29 biopsy from a vehicle control scar. For the analysis, a number of epidermal and dermal variables were quantified, including parameters measured from 4 equally spaced depth quartiles. **B)** Plots of mean separation of (αCT1 treatment minus Vehicle control averages) of intensity of eosin staining (% Eosin top), cell nuclei density (middle) and blood vessel density (bottom) in the 4 dermal quartiles. The only variable measured from scar epidermis and dermis that showed a significant treatment effect (p<0.05 – red text) was the intensity eosin staining, a measurement related to the collagen content of the tissue.

### αCT1 Prompts Changes in the Collagen Organization of Human Cutaneous Scar Tissue

Based on the indication from H&E staining that αCT1 may have effects on scar ECM (Fig. 2B), we undertook a rigorous analysis of the ECM in scar biopsies of all patients in the Phase I trial. Sections of the dermal scar biopsies collected at day 29 were stained with Picrosirius (PS) red to enhance birefringence of collagen fibers and full-width images of scar sections were scan-montaged on an automated scanning microscope using a 20x objective (Fig. 3). Initially this was achieved using a single, circularly polarized angle of light, but as discussed below, for quantification purposes we later devised an approach based on imaging at multiple polarization angles. Vehicle control and 100 μM αCT1-treated scar samples from the same patient in cohort 3 at 29 days are shown in Figures 3A and B, respectively. The most conspicuous difference between the two scars from the same patient was greater alignment of collagen bundles in the control scar relative to the αCT1-treated tissue (Figs. 3C & D). Also notable in the αCT1-treated scar were whorls of randomly oriented collagen bundles reminiscent of those in unwounded skin (Figs. 3D). These differences are emphasized in Figures 3E-G, where angular disposition of collagen bundles from PS red birefringence has been color-coded over 180 degrees. The increased heterogeneity of organization in the αCT1-treated scar is showcased by the broader and more variable palette of colors, reflecting the more random arrangements of collagen bundles relative to the vehicle control.

**Figure 3.**
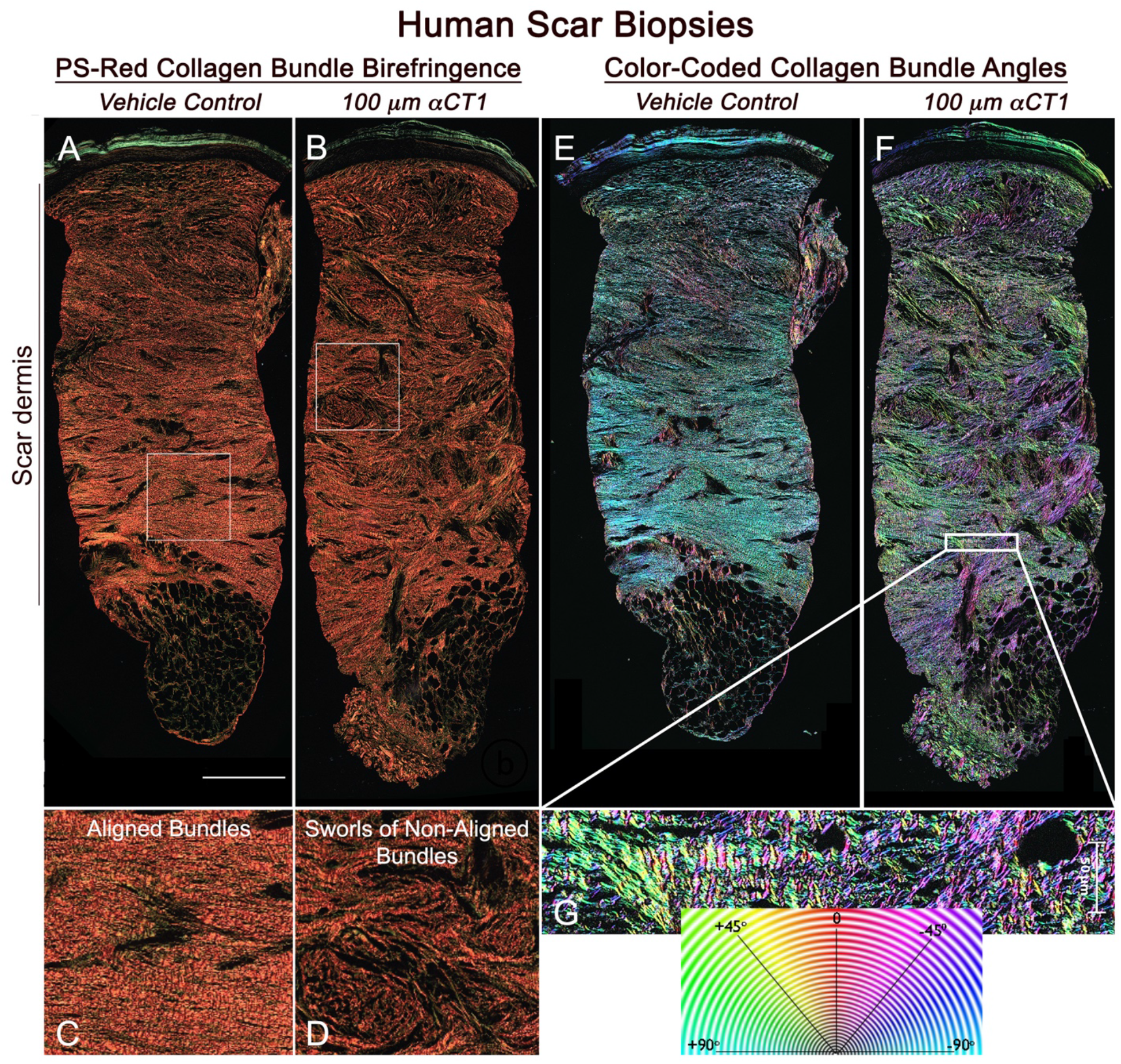
αCT1 effects on collagen organization in the human 29-day scar biopsies. **A)** Olympus VS120 scan (20x objective) of a vehicle-control scar biopsied from the left arm of an individual in cohort 3 - i.e., the patient group receiving the 100 μM therapeutic dose of αCT1. **B)** Corresponding scan of collagen birefringence of the αCT1 treated scar on the right arm of the same patient. **C and D)** Higher magnification views of the organization of collagen bundles in the boxed regions of (A) and (B), respectively. **E and F**) Collagen bundle angles in full-width sections of scars shown in (A) and (B) are color-coded over 180 degrees using OrientationJ software. H) Higher magnification views of boxed region in (F). Collagen bundles show a high degree of alignment in the control scar. In the αCT1-treated scar collagen bundles tend to be more randomly arranged and form larger sworls reminiscent of structures seen in unwounded skin. Scale = 0.5 mm

Next, we used MatFiber^32^ and the Circular Statistics Toolbox^33^, analytical tools developed within Matlab^34^, to calculate metrics of the organization of scar ECM. The first of these was the circular variance of the alignment angle of bundles of collagen as an average within 20 depth sub-regions into the scar dermis. Circular variance provides a measure of the breadth of collagen bundle angles; a dermal sub-region with all bundles aligned anisotropically in the same direction will give a circular variance of zero, whereas totally randomly aligned collagen bundles will give a circular variance approaching one^33^. While circular variance is a unitless measure of angle variance, variance in density across the measured sub-regions can bias the calculated value, so circular variance was weighted by density to correct for bias. This metric is subsequently referred to as bundle disorganization. Additionally, we calculated variance in bundle disorganization within depth sub-regions to provide an index of regional heterogeneities in fiber organization, e.g., bundle whorls. We refer to this measurement as bundle disorganization variance. To ensure that we obtained a complete array of collagen fibers present in each sample, the sections were serially imaged at six polarization angles - 0°, 15°, 30°, 45°, 60°, & 75° using customized electronics and software that we engineered for this purpose on an automated Olympus VS120 microscope (Supplemental Fig. 2). This enabled our analyses to be based on multi-angle images of circularly polarized light, providing increased rigor to quantification of collagen bundle organization. Density of collagen bundles per unit tissue area was also measured from the scan-montaged fields of scar.

No significant effects were found for any of the 3 parameters measured in the cohorts receiving the lowest (cohort 1: 20 μM) and highest (cohort 4: 200 μM) doses of αCT1 (Fig. 4). Overall collagen bundle disorganization, disorganization variance, and density were all slightly higher in αCT1-treated than control wounds in cohort 1, but these effects were not significant (Fig. 4A). The cohort receiving 50% of the therapeutic dose (cohort 2: 50 μM αCT1) had a significant treatment-by-depth interaction effect on bundle disorganization (p<0.05), but no significance overall (Fig. 4A, top graph). For patients receiving the therapeutic 100 μM dosage of αCT1 (cohort 3), both treatment overall and treatment-by-depth interaction effects on bundle disorganization were significant (Figs. 4A & B, top graphs; p<0.0076). Bundle organization variance also significantly varied by treatment overall and by interaction with dermal depth in cohort 3 (Fig. 4A and B; middle graphs, p=0.0097). There were no significant αCT1-treatment associated effects on collagen density in any of the four patient cohorts (Figs. 4A & B, bottom graphs) – though cohort 3 showed a trend toward peptide-treated wounds being slightly more dense than control scars (p=0.0666).

**Figure 4.**
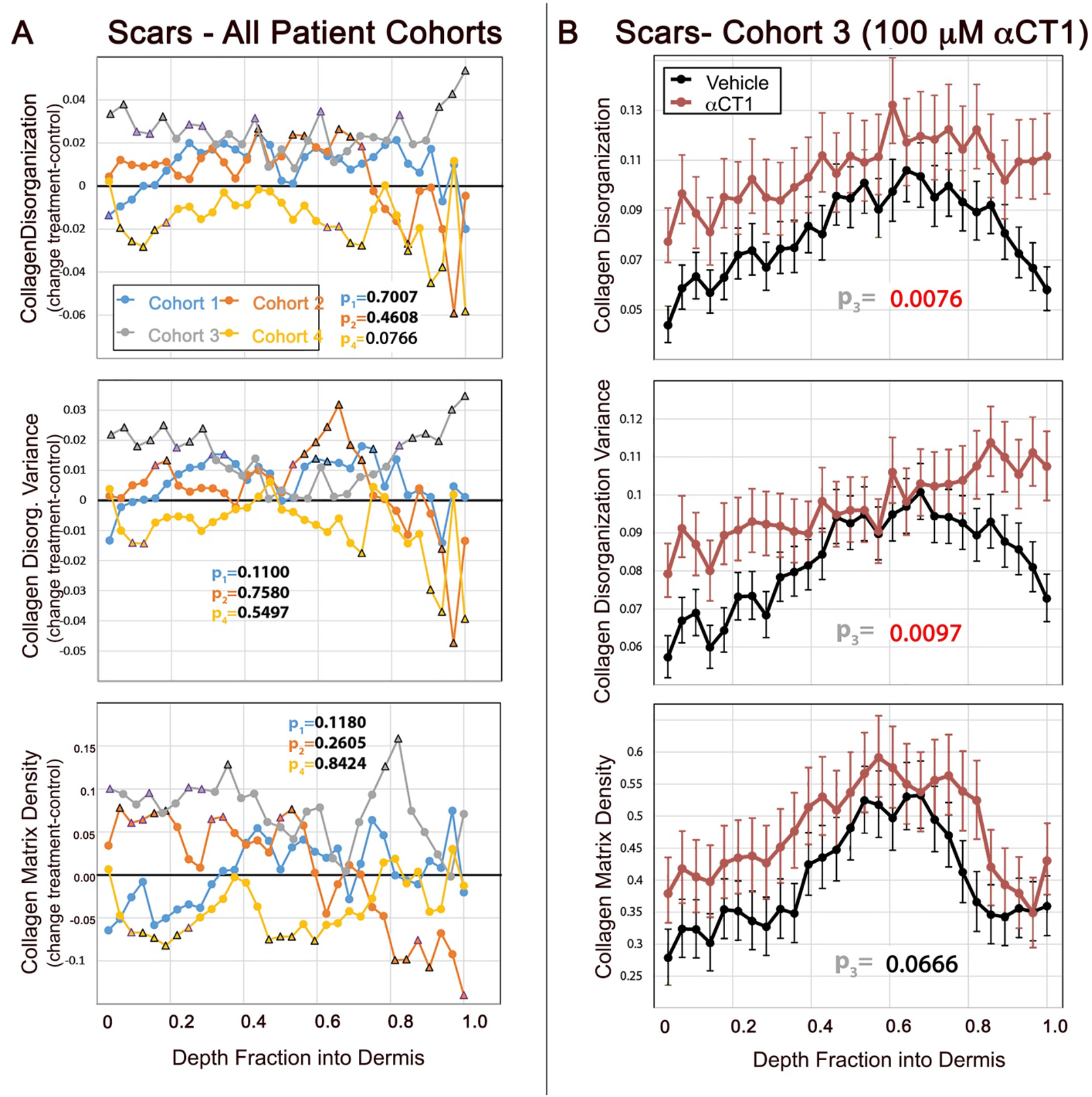
Quantification of αCT1 effects on collagen organization in the human 29-day scar biopsies. **A)** Differences between mean treatment and control measurements for collagen bundle density-adjusted circular variance (disorganization), variance in collagen bundle disorganization, and collagen fiber density from the Phase I clinical trial. Plots are shown for each dosage cohort from the most superficial part of the dermis (right) to the deepest part of the dermis (left). The measurements were made on images taken at multiple polarization angles. Black triangles on plots indicate a statistically significant difference (p<0.05) between treatment and vehicle at individual fractions of depth into the dermis. Trending points (0.05≤p<0.08) are denoted with a purple triangle. **B)** Cohort 3 collagen bundle disorganization, disorganization variance, and density by depth into dermis for αCT1-treated (red) and vehicle control (black) scars imaged at multiple polarization angles. Error bars ± standard error (SE). Treatment and control measurements of dermal collagen disorganization and its variance differ significantly in the cohort receiving the therapeutic 100 μM dose of αCT1 - red p values on plots.

### αCT1 Causes Changes to Collagen in Rat and Guinea Pig Scars Similar to those in Human

Significant differences in collagen organization were observed between treated and control scars sampled from within the 22 patients in cohorts 2 and 3 – comprising individuals subject to 50 μM or 100 μM of αCT1. To further validate these results, and also to explore αCT1 mechanism-of-action, additional studies using translationally relevant wound healing animal models were conducted. We used the Sprague Dawley (SD) rat and the Institue of Armand Frappier (IAF) hairless guinea pig for this purpose, the latter of which have skin of similar thickness and structure to humans, including shallow dermal papillae and small, vellus hair follicles.^35^ The experimental design and end-points employed in these animal studies purposely recapitulated key aspects of the human clinical trial (Supplemental Fig. 3A). Specifically, within-subject comparison of the effects of acute treatment with the therapeutic 100 μM dose of an αCT1 gel relative to a vehicle control gel was carried out and assessed based on multi-angle birefringence of PS-red stained scar samples in a randomized, blinded manner. One key aspect of our animal model approach was the generation of a mechanical environment during wound healing in the loose-skinned small animal models that more closely resembled that of tight-skinned humans. To maintain a higher mechanical tension in the skin, as is naturally present in humans, we secured 0.8 mm thick circular silicon splints around wounds to exert uniform tension on the wound area over the course of the study (Supplemental Fig. 3B). Additionally, instead of sampling at a single time point post-injury, as was performed in the Phase I trial, rat and guinea pig scars were biopsied at 2-, 4-, and 6-weeks following treatment, enabling the time course of changes to collagen organization in healing injuries to be followed.

The effects of 100 μM αCT1 on collagen alignment in rat skin showed effects resembling those observed in the human clinical trial samples (Fig. 5), wherein αCT1-treated scars had consistently higher levels of collagen bundle disorganization (Figs. 5A & B) and disorganization variance (Figs. 5C & D) across dermal depths. However, though within-subject treatment effects reached significance for disorganization and disorganization variance at 2 and 4 weeks respectively, the overall magnitudes were less than those in humans receiving the therapeutic dose. Interestingly, we also noted that whilst collagen disorganization and variance differed significantly between treatment and control throughout the dermis in scars sampled at week 2 (Fig. 5A), the differences were only maintained in the upper part of dermis (i.e., nearer the epidermis) in the 4-week sample (Fig. 5B). The collagen density of αCT1-treated rat scars was significantly increased at week 2 (Fig. 5E), but at week 4 the depth profile of density was not significantly different between treated and control scars (Fig. 5F). These data, along with the trend observed in the human scars (Fig. 4), suggests that αCT1 transiently increases the density of collagen in the scar. Rat scars were also biopsied at week 6, however, these were excluded from further analysis, as due to the successful efforts of a number of our more dexterous and enterprising rats splint loosening became a persistent issue towards the end of the 6-week study period.

**Figure 5.**
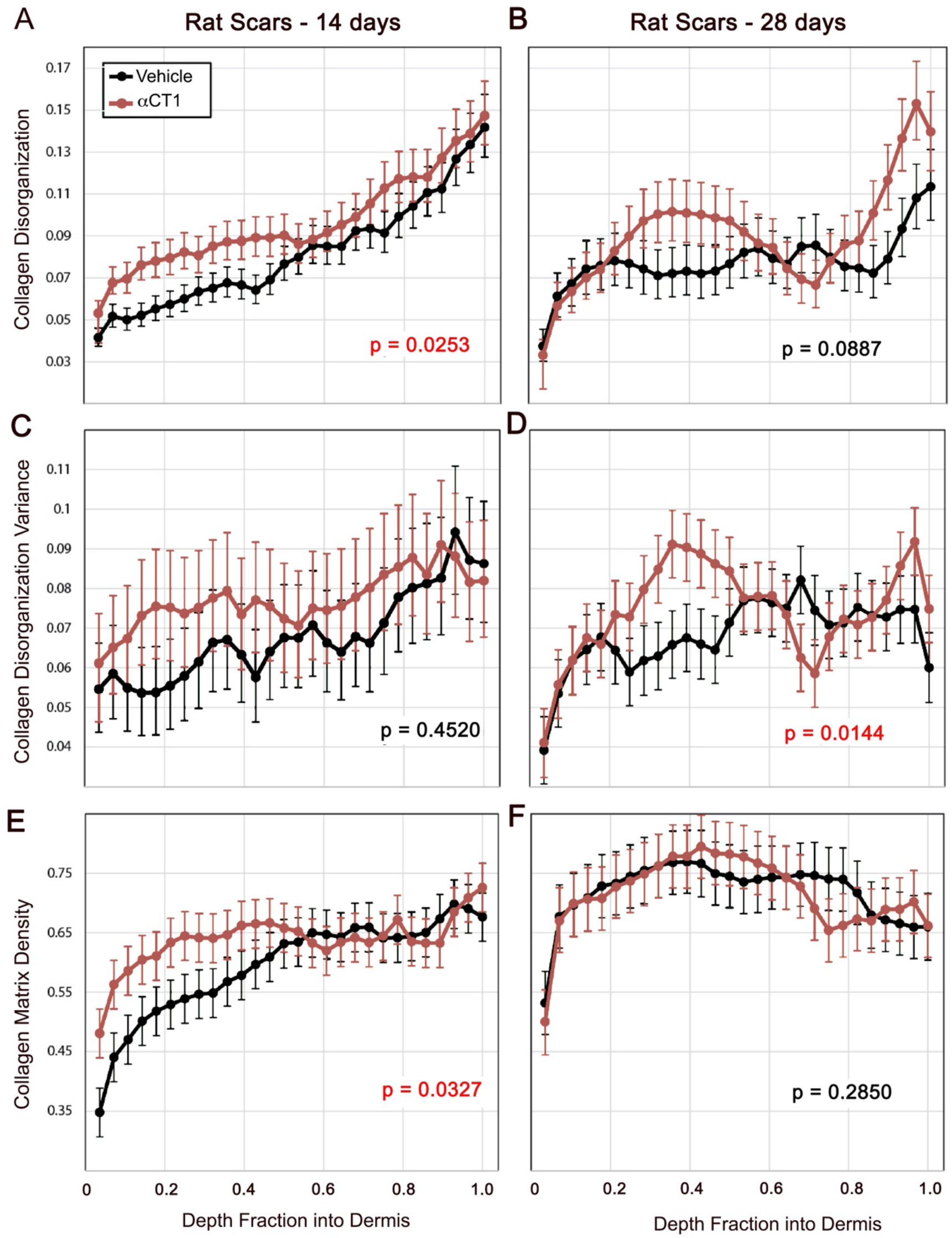
Quantification of αCT1 effects on collagen bundle organization in rat cutaneous scar biopsies 14- and 28-days post-injury. Collagen bundle disorganization, disorganization variance, and density by depth into rat dermis for for αCT1-treated (red) and vehicle control (black) scars imaged at multiple polarization angles 14 and 28 days following cutaneous wounding and splinting of those injuries. Error bars ± standard error (SE). Red p values on plots indicate significant (p<0.05) overall differences between treatment and control scars.

Guinea pigs provided larger treatment effects, retained their skin splints more consistently through to the end of the study period, and overall modeled the human Phase I trends more closely than was achieved with rats (Fig. 6). Collagen bundle disorganization in αCT1-treated scars was elevated over that of controls at all three time points, showing notably large and significant increases at weeks 4 and 6 (Fig. 6A). Disorganization variance showed a similar pattern with the difference between treated and vehicle control scars becoming progressively larger over the time course of the study (Fig. 6B). The strength of the effect of αCT1 treatment on bundle disorganization variance was dependent upon depth into the tissue, with a diminished effect in the superficial and most basal parts of the dermis. Collagen density at week 4 was significantly higher in αCT1 treatment samples, but was not different from control at weeks 2 and 6 (Fig. 6C). This transient increase in collagen density was similar to that observed in rat and human samples, although more pronounced in significance. Overall, the results from all 3 species suggest that αCT1 accelerates the differentiation of scar ECM, causing early transient increases in collagen density without increasing the fibrosis of the final long-term scar.

**Figure 6.**
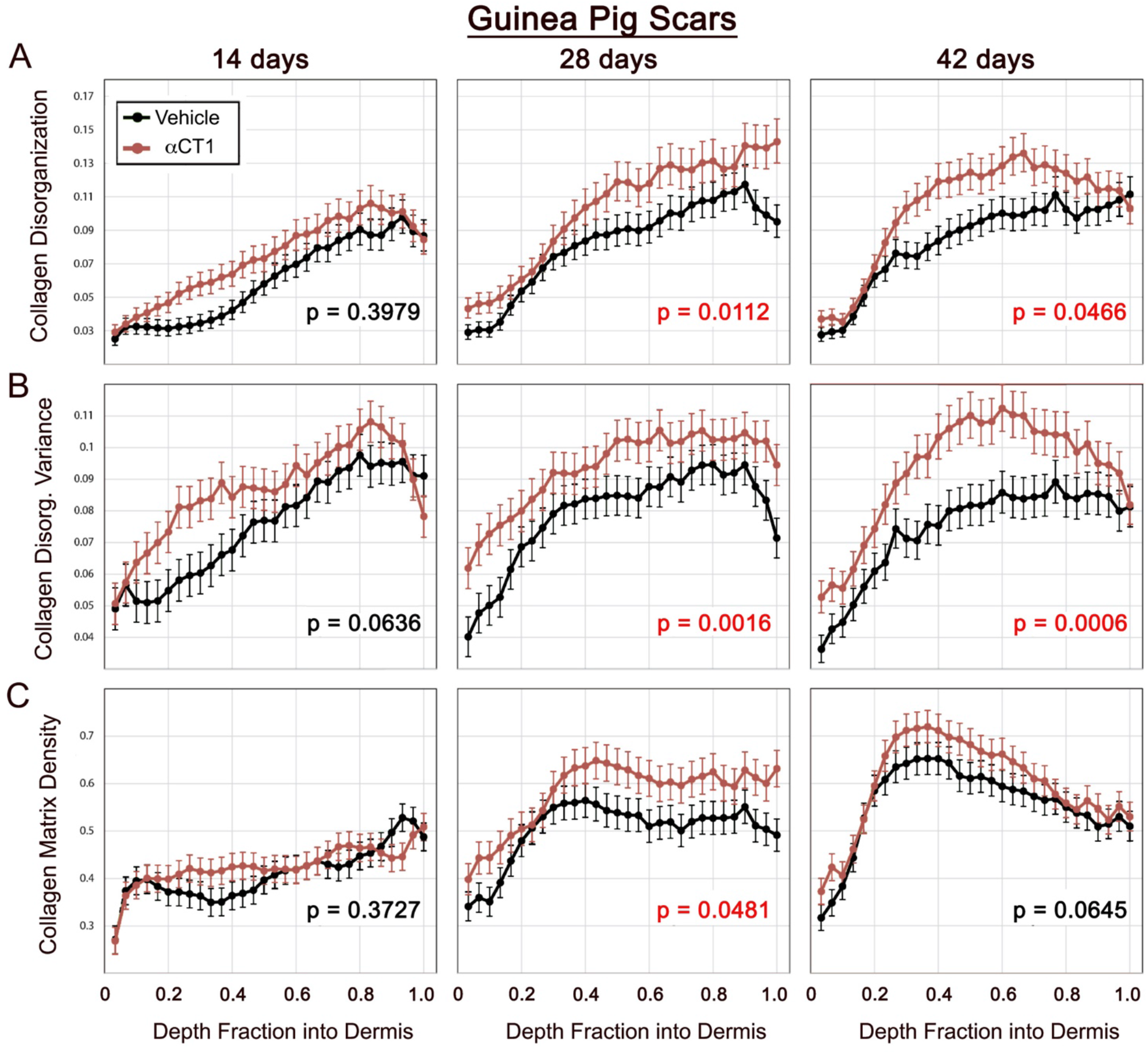
Quantification of time course of αCT1 effects on collagen organization in cutaneous scar biopsies from guinea pig. Collagen bundle disorganization, disorganization variance, and density by depth into guinea pig dermis for αCT1-treated (red) and vehicle control (black) scars imaged at multiple polarization angles 14, 28 and 42 days following cutaneous wounding and splinting of those injuries. Error bars ± standard error (SE). Red p values on plots indicate significant (p<0.05) overall differences between treatment and control scars.

### αCT1 Prompts Changes in Rate and Directionality of Fibroblast Motility

The migratory behavior of fibroblasts, the main ECM-protein-expressing cell-type, canonically strongly influences scar organization.^36–40^ To explore the possibility that the changes in scar structure we observed in response to αCT1 were influenced by alterations in cell motility, we undertook experiments on fibroblasts *in vitro*. As a first step, we surveyed whether fibroblast dynamics altered in response to αCT1 in a standard cell-cultured scratch-wound assay (Fig. 7). Fibroblasts were grown to sub-confluence for 24 hours, treated with different doses of αCT1 (1-180 μM), control peptide or vehicle control for a further 24 hours. The monolayers were then scratch-wounded with a 200 μL pipette tip and video recorded over the subsequent 6 hours. Measurements from these videos indicated that αCT1 prompted a dose-dependent increase in the relative rate of migration of individual cells back into the scratch wound (Fig. 7A).

**Figure 7.**
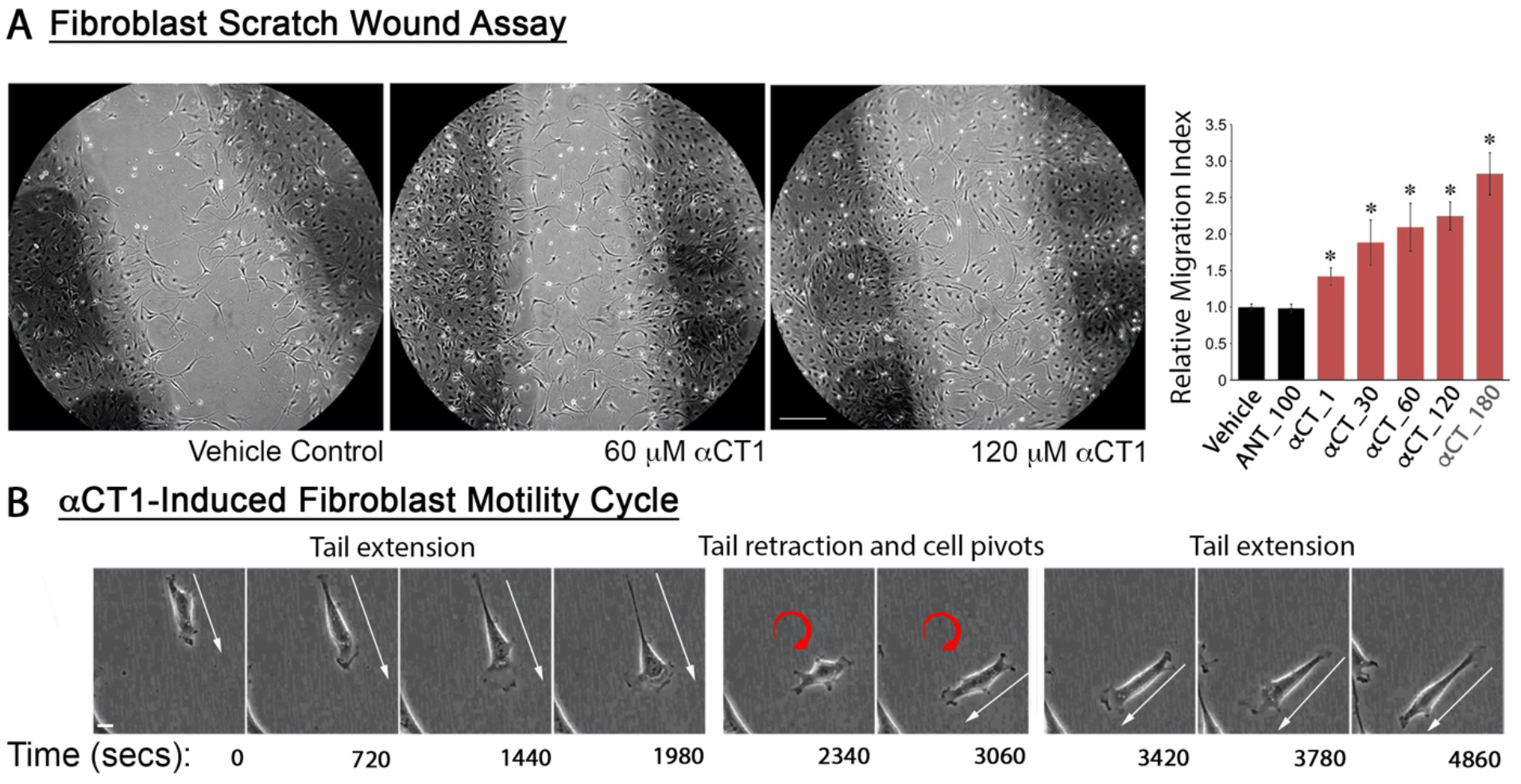
αCT-1 effects on the rate and directionality of fibroblast migration. A) Phase-contrast images of (10x objective) of NIH-3T3 fibroblasts in a scratch-wound migration assay 24hrs after scratching and exposure to Vehicle control, 60μM αCT-1, and 180μM αCT-1. The scratch edge is demarcated by the darkly colored strips within the circular area. Graph in (A) shows fibroblast migration was quantified by counting the number of cells in 10 fields both inside and outside the wounded area. The “Relative Migration Index” is defined as vehicle control-normalized ratios of cells inside to cells outside the scratch area in the different treatment and control conditions. * p < 0.002 αCT1 versus vehicle and antennapedia peptide controls, N=5 experimental replicates. **B)** Sequence of images from supplemental movie 1 of a fibroblast (initial position asterisked in video) in a scratch-wound migration assay 24hrs after exposure to 180 μM αCT1. The cycle of behavior shown involving cellular tail extension, release and pivoting is frequently exhibited by fibroblasts exposed to αCT1, resulting in reduced directionality of motion. Scale A = 100 μm; B= 5 μm.

We next focused on the behavior of individual cells during migration into the scratch wound. Video sequences revealed distinct cellular differences between “wounded” cultures receiving αCT1 and control solutions (Fig. 7B; Supplemental videos 1 and 2). In response to αCT1, fibroblasts appeared more elongated and active than cells in control cultures. We also noted that treated fibroblasts changed direction more frequently - with cells appearing to take more random (less linear) paths across the substrate in the presence of αCT1. These treatment-associated changes in cellular direction followed a choreographed sequence that rarely occurred in control conditions - illustrated in figure 7B. During forward progression in an αCT1-treated culture, a fibroblast is seen forming an extended tail. As the tail reaches a maximum extension, the adherent base of the tail detaches and rapidly propagates forward toward the body of the cell. The apparent recoil of the formerly extended tail coincides with a pivot and change in direction by the fibroblast. The random paths taken by cells in αCT1 cultures seemed to result from repetition of this process of tail extension, release and pivot.

To further investigate the pivoting behavior and determine the extent to which it was a dose-dependent effect of αCT1, we set up new cultures of fibroblasts and video-recorded the cells over eight hours of migration under six different experimental conditions: αCT1 dosages of 10, 50 and 100 μM, a 100 μM dose of inactive αCT1 peptide control, a 100 μM dose antennapedia peptide control and a no treatment control. A custom suite of software that we developed was used to measure fibroblast dynamics and model its consequences.^41^ Quantitative analysis of fibroblast motility revealed that treatment with αCT1 caused a dose-dependent decrease in the directional persistence of cells, more random migration tracks (Fig. 8A) and a flatter distribution of directional angle changes, averaged amongst all cells at all time steps (Fig. 8B). To quantify the central tendency of cell directional persistence, we calculated the mean vector length (MVL) of the angle change distribution (Fig. 8C - a measure of alignment around a central angle), as well as the number of angle changes within +/- 30° of the previous cell orientation (Fig. 8D). Both alignment measures demonstrated a dose-dependent decrease in the directional persistence of cells in response to αCT1 relative to control conditions (Figs. 8C & D).

**Figure 8:**
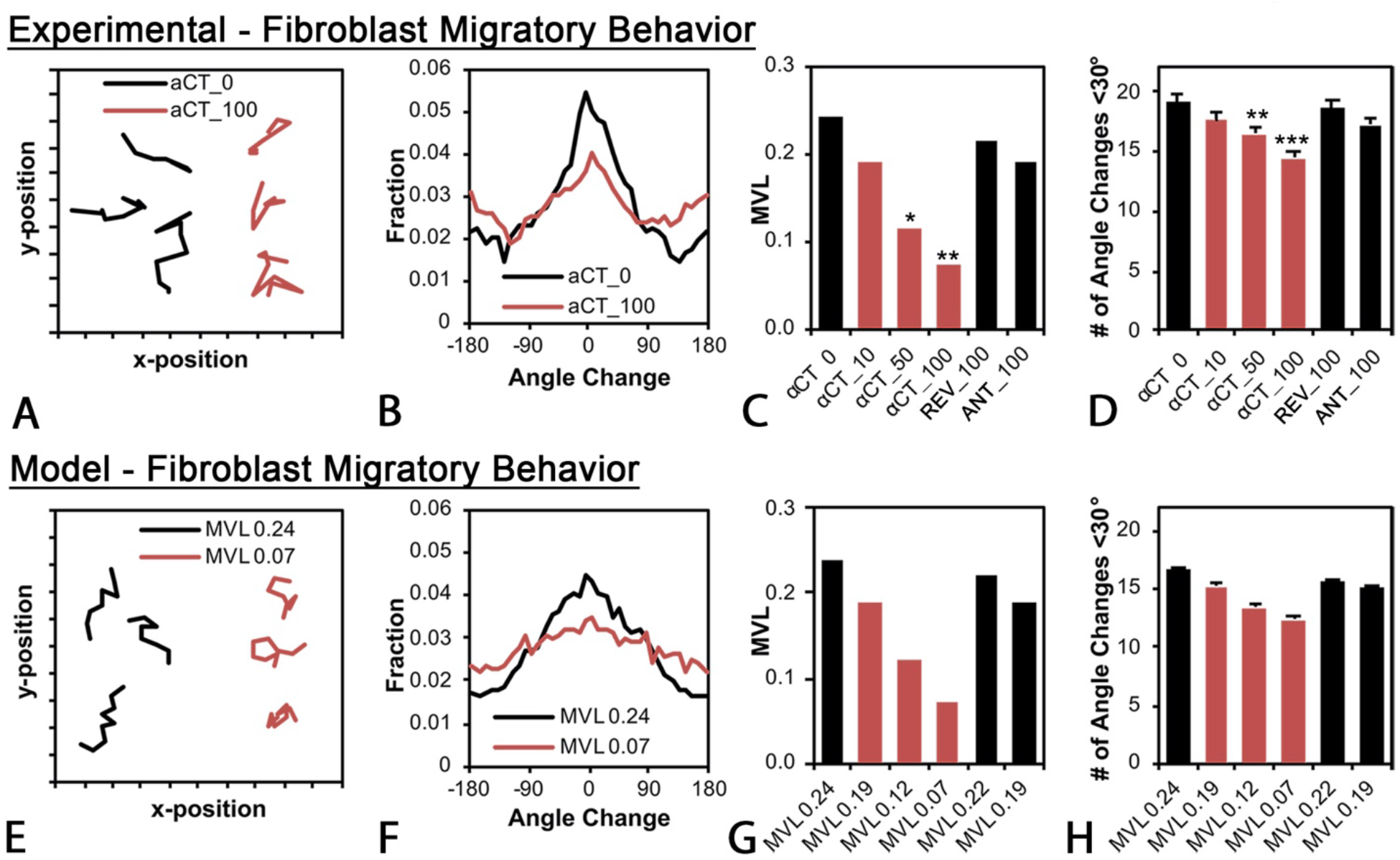
Experimental (A-D) and agent-based modeling (E-H) effects of αCT1 on fibroblast migration patterns. **A)** Migration paths of NIH 3T3 fibroblasts tracked over eight hours in experimental 100 mM αCT1 treatment (αCT_100, brown lines) or no treatment control (αCT_0, black lines) conditions. A total of 911 cells (200-250 per group) were tracked in N=3 experimental replicates. **B)** Distribution in average directional angle changes (pivots) of cells in 100 μM αCT1 (αCT_100, brown lines) treatment or no treatment control (αCT_0, black lines) conditions, taken as means of all cells and all time steps. **C)** Mean vector length (MVL) of the angle change distribution for cells receiving 10 (αCT_10), 50 (αCT_50), or 100 μM (αCT_100) αCT1, 100 μM inactive control peptide (REV_100), 100 μM antennapedia control peptide or no treatment control (αCT_0). **D)** Number of angle changes within +/- 30° of the previous cell orientation for the experimental groups in (C). Both (C) and (D) demonstrate a dose-dependent decrease in the directional persistence of cells. The effects of αCT1 were simulated *in silico* using a computational agent-based model. To simulate treated and control cell behaviors, directional persistence was set to levels equivalent to values observed in the experiments for the differing conditions (i.e. values from C). **E)** Simulations of fibroblast migration paths in peptide treatment (brown lines) and control (black lines) conditions. **F)** Modeled distribution of average directional angle changes of cells in simulated αCT1 (MVL0.07-brown) and control (MVL0.24 -black) conditions. **G)** Simulated mean vector length (MVL) of the angle change distribution for cells receiving 10 (MVL0.19), 50 (MVL0.12), or 100 μM (MVL0.07 MVL0.24) αCT1, 100 μM inactive control peptide (MVL0.22), 100 μM antennapedia control peptide (MVL0.19) or no treatment control (MVL0.24). **H)** Number of angle changes within +/- 30° of the previous cell orientation for the modeled groups in (G). Bars D and H = standard deviation. ns=not significant, *=p <0.1,= **=p <0.05 and ***=p<0.0001 compared to groups receiving no treatment (αCT1_0).

### Effect of αCT on scar collagen structure in computational simulations

In addition to providing measurements of cellular motility, our computational suite can simulate the effects of treatments in an agent-based model of cell-matrix interactions. This model can be parameterized to reproduce the behavior of cell populations at microscopic scales, and at the macroscopic tissue level, predict the structure of scars that might result from the collective actions of cellular agents. ^41^ To simulate fibroblasts, we set a directional persistence adjustment factor in the model equivalent to the MVL values observed in the experiments (i.e., Fig. 8C). We determined that modeled cell migration paths, closely agreed with experimental findings showing αCT1 produced more random (linear) migration tracks (Fig. 8E), a flatter distribution of directional angle changes (Fig. 8F), dose-dependent decreases in the angle change alignment (MVL, [Fig. 8G]), and dosedependent decreases in the number of angle changes within +/- 30° of the previous cell orientation (Fig. 8H).

To test the effect of αCT1 treatment in our computational model at the tissue-scale, we simulated 28 days of healing time with ~9000 cells under four different conditions: 0% mechanical strain without αCT1, 0% strain with αCT, 2.5% strain without αCT1, and 2.5% strain with αCT1 (Fig. 9). Parameter settings on the latter two conditions sought to mimic the mechanical environment that occurs in a human skin wound and in the splinted wounds that we used in the animal experiments. Without mechanical strain and without αCT1, the model predicted disorganized collagen bundle orientation within the simulated scar tissue, with the exception of small alignment zones at the interface of the wound and normal tissue regions (Fig. 9, top left panel). In the presence of uniaxial mechanical strain without αCT1, the model predicted highly organized collagen orientations within the scar tissue, which was produced as cells oriented parallel to the uniaxial tension (horizontal, x-direction) and then deposited new collagen parallel to their own orientation (Fig. 9, top right panel). These simulated matrices matched the patterns of collagen bundles observed in vehicle control scars. We simulated the effect of αCT1 by turning down a directional persistence adjustment factor. This resulted in low cell alignment with directional cues and low levels of bundle organization for both 0% strain and 2.5% strain (Figs. 9 bottom left and right panels, respectively). The simulated matrix for treated wounds under tension strongly resembled the scar histoarchitecture observed for wounds receiving the therapeutic dose of αCT1 in the Phase I human biopsies, as well as in the animal studies. Thus, the predictions from the agent-based model simulations agreed with the experimental findings and are consistent with the hypothesis that αCT1 effects on directional persistence of fibroblasts can compound over a system of 9000 cells to potentially explain the collagen bundle disorganization patterns that were observed *in vivo*.

**Figure 9.**
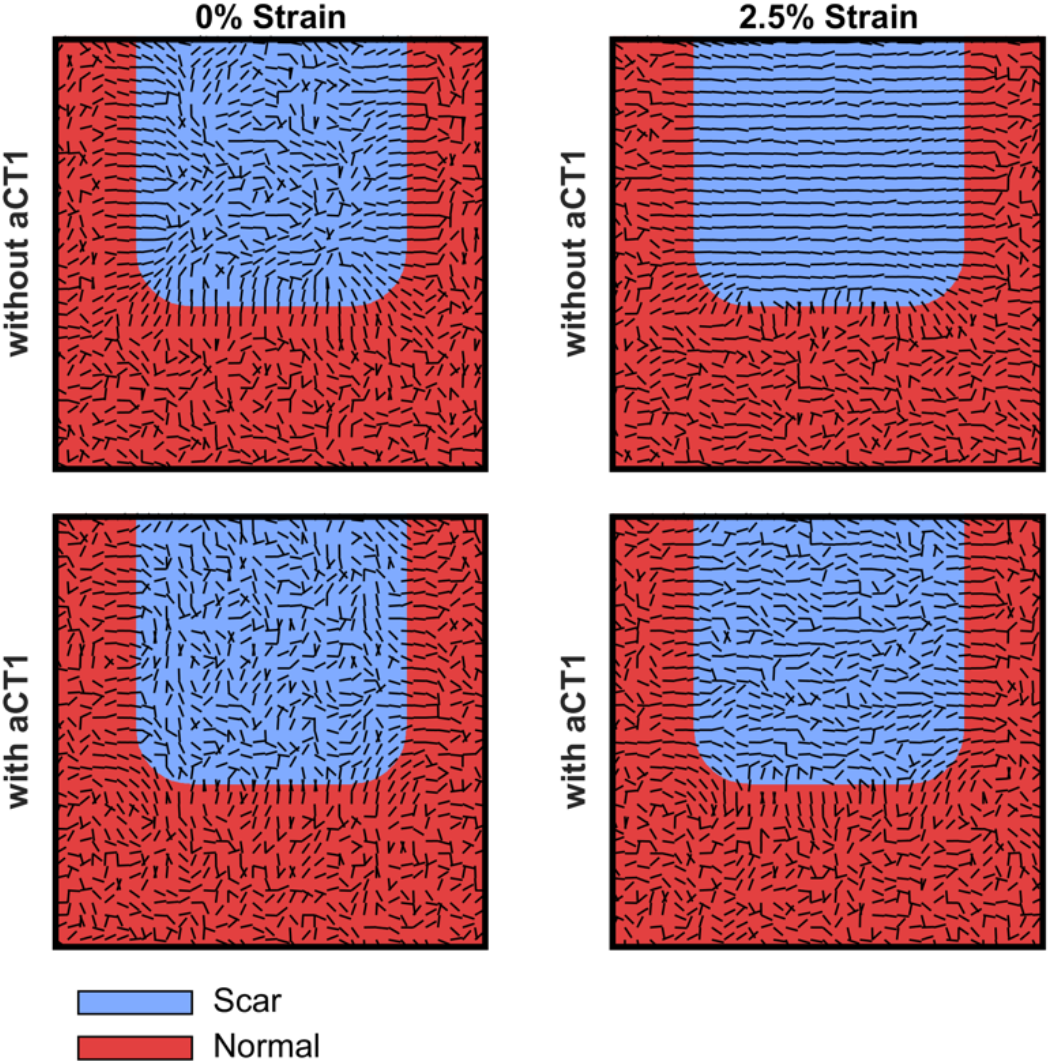
Computationally simulated scar structures predicted from agent-based modeling of fibroblast motility in presence or absence of αCT1. The model simulated 28 days of healing time with ~9000 cells under four different conditions: 0% mechanical strain without (top left hand scar simulation) and with (top right hand scar simulation) αCT1 and 2.5% strain without (top right hand scar simulation) and with (bottom right hand scar simulation) αCT1. The latter two conditions modeled vehicle control and peptide treatment conditions from the Phase I clinical trial. MVL values with and without αCT1 were set to those measured for the therapeutic 100 μM dose of the peptide or no peptide treatment, respectively. Collagen bundle orientation within the simulated scar tissue was disorganized in all cases, except in the presence of mechanical strain without αCT1 – as is seen in normal cutaneous scarring in humans, and as exemplified by vehicle control scars from the Phase I clinical trial. The computational simulation suggests that αCT1 desensitizes cells to mechanically based alignment cues causing more random migration paths, which in turn results in the generation of disorganized collagen matrices within the scar.

## Discussion

In this study, we show that topical treatment with the Cx43 CT mimetic peptide αCT1 in the first 24 hours after skin wounding affects the organization of collagen bundles in dermal granulation tissue formed in the subsequent weeks of healing. This effect was observed in human skin, as well as in that of rat and hairless guinea pig wound models. Collagen bundles in treated scars were more randomly aligned and their overall three-dimensional (3D) organization resembled that of normal, unwounded skin. Moreover, we observed that cultured fibroblasts treated with αCT1 showed dose-dependent decreases in directionality of movement, resulting in increased randomness in the migration paths taken by the cells. An agent-based model parameterized with motility data from fibroblasts predicted collagen organizations in computationally simulated scars consistent with those observed from human clinical testing and animal experiments. In sum, our study indicates that the mode-of-action of αCT1 in mitigating cutaneous scarring may involve inducing patterns of collagen organization that quantitatively resemble those found in normal unwounded skin, potentially via effects of the peptide on fibroblast motility.

The histology of unwounded skin and cutaneous scar in control subjects from the Phase I clinical trial agree with reports in the literature.^42–45^ The tendency of collagen bundles to be highly aligned in normal dermal scars has been proposed as a reason as to why scar tissue has increased susceptibility to mechanical disruption and distinguishes itself visually from surrounding unwounded skin.^46–48^ Our survey of the histological characteristics of epidermis and dermis of αCT1-treated scars further reinforces the latter point. The only consistently significant treatment-associated change detected from the variables examined was on the organization of collagen bundles in the dermis. It is unclear why this alteration to dermal ECM structure might make scars stand-out less from adjacent skin, though it may relate to differences in how light is refracted by aligned versus more isotropically arrayed networks of collagen – as has been suggested from studies of corneal wounds.^49^ Applied mechanical strain on remodeling porcine scars has been reported to result in dose-dependent increases in collagen alignment and this experimental model suggests itself as an approach to further investigate how differing levels of collagen disorganization could mechanistically relate to the visual appearance of scars.^50^

The impetus for this further investigation of αCT1 effects on cutaneous scarring came from a Phase II trial evaluating its effect on treatment of surgical wounds, which showed that the peptide mediated clinically meaningful levels of scar reduction.^31^ Acute external application of an αCT1-containing gel (within the first 24 hours) onto one of a pair of 1 cm laparoscopic surgical incisions improved longterm scar appearance relative to the untreated scar, as confirmed both in blinded appraisals by clinicians and self-assessment by patients. The emergence of the treatment effect was gradual, with incisions to which 100 μM αCT1 was administered showing a 13% improvement over paired controls at 3 months, 17% improvement at 6 months, and 47% improvement at 9 months. Interestingly, blinded assessment also indicated that there was no appreciable difference in αCT1-treated and control scars at the 1-month time point. Although the improved scarring outcome associated with αCT1 at 9 months could not have been predicted by visual inspection of incision locations 1 month after surgery, we hypothesized that scar histoarchitecture at this early stage may have been altered by treatment in a manner that was not qualitatively obvious from the surface of the healed wound. The results of this study support this hypothesis.

Our data suggest the 3D order of dermal collagen bundles established in the first 2 weeks of scar formation may provide a framework that is critical to the long-term structure and function of the fibrotic repair. Results from earlier experiments on mouse and pig skin wounds presaged this conclusion. In these preclinical studies it was shown that acute αCT1 treatment increased wound closure rate, decreased inflammatory neutrophil infiltration, and reduced granulation tissue area.^29, 30^ Of particular relevance, it was also determined that αCT1 treatment was associated with scars with improved mechanical properties (stress and strain following extension to tissue rupture) three months but not one month after injury.^29^ Together with the results of present study, this data supports a conclusion that αCT1 alters the differentiation of granulation tissue, eventually manifesting as a fibrotic repair with improved mechanical and visual properties after long-term scar remodeling.

A key question relates to how acute αCT1 treatment mediates this progressive long-term effect on scar remodeling. Earlier work indicated that following topical application *in vivo*, αCT1 cannot be detected at injuries beyond 48 hours.^29, 51^ Thus, it is unlikely that the peptide is active in the healing wound beyond the first few days post-injury and certainly would not be present in any amount at 9 months, when effects on scar appearance were most pronounced.^31^ One potential source of insight here may come from how ECM alignment quantitatively varied with depth in the scars. In all three species (i.e., Figs. 4–6), the more basally located collagen in the dermis consistently showed significantly increased levels of disorganization and alignment variance compared to bundles located closer to the epidermis. These trends with depth were enhanced by αCT1 treatment and in guinea pigs were evident by 2 weeks (Fig. 6). The most basally located collagen is amongst the first laid down by fibroblasts^52^ - and again, our observations are consistent with the concept that ECM, from the earliest stages of granulation, may provide a 3D template upon which subsequent scar order is constructed over ensuing weeks and months. That the strongest effects of αCT1 were found in the deepest part of the dermis, may also partly explain the delayed skin surface appearance improvements seen in Phase II clinical trials at 9 months. Of interest in ongoing studies are questions concerning how αCT1 might influence the structure and molecular composition of the provisional wound matrix formed sub-acutely during the first week following cutaneous injury,^53, 54^ as well as effects of the peptide on differential expression of collagen isoforms during early wound healing. Type III collagen typically predominates over collagen I during initial granulation tissue formation, and increases in the ratio of collagen III to I in the wound space have been linked to improved scar outcomes in fetal skin^55–57^ and the acomys regenerative skin mouse.^58^

Furthermore, the data from the present study indicates that αCT1 effects on scar organization likely derived from peptide-induced alterations of fibroblast migration patterns. Mathematical modeling, as well as experimental work, has strongly implicated directionality in the movement of collagen-secreting cells as critical to the differentiation of scar ECM.^36–40^ These studies includes our own computational simulations, which have elucidated the key role played by gradients of chemokines and mechanical strain in guiding fibroblast homing and the subsequent alignment of collagen bundles laid down by these cells.^41^ The models in the present study indicate that the sensitivity of fibroblasts to αCT1 may be strongly influenced by mechanical strain. Specifically, the simulations suggest that the peptide renders cells insensitive to biomechanical inputs during scar differentiation. The results from splinted wounds align with this hypothesis. Ongoing work would usefully address whether the prediction from modeling is born out in experimentally.

The most notable change in the fibroblast motility caused by αCT1 was the increased frequency of the behavior illustrated in Figure 7B – wherein during movement cells showed a tendency to extend unusually long trailing edges (tails), which were then released and retracted in association with a pivot-like change in the direction of the cell. In the presence of the peptide, fibroblasts appear to repeat this behavior in cycles, culminating in increasingly random migration paths. The reorientation and directed migration of fibroblasts spurred by mechanical strains requires the proper functioning of various integrins and focal adhesion kinase (FAK) at focal adhesions contacting the extracellular matrix.^59–63^ Focal adhesions at stress fiber termini in the cytoplasm are known to be critical to maintenance of anisotropic movement by fibroblasts.^63^ Following tail retraction, trailing adhesions are actively regenerated and associated stress fibers are remodeled; with tensile forces generated by actin networks consequently redirected according to trailing adhesion location. The long tail extensions and delayed release of terminal adhesions in treated cells suggests that αCT1 may interfere with dynamic processes regulating trailing focal adhesion turnover. Although the molecular basis of this fibroblast behavior remains to be determined, previous studies in migrating endothelial cells have indicated that αCT1-mediated disruption of Cx43/ZO-1 complexes resulted in collapse of the organized F-actin cytoskeleton and the appearance of actin nodes.^64^ Cx43 has also been shown to influence migration directionality via effects on microtubule dynamics^65^ and is key to directed migration of glia in the cerebral cortex^66^ and neural crest cells to the developing heart.^67^

In addition to published clinical trials on scar mitigation,^31^ we have reported outcomes of Phase II testing of αCT1 in the treatment of diabetic foot (DFU) and venous leg (VLU) ulcers. The peptide’s demonstrated efficacy in increasing the rate of resolution of both types of pathological wounds.^68, 69^ It is unknown whether the mode-of-action of αCT1 in prompting accelerated closure of chronic wounds is related to the mechanism presented here by which the peptide may influence healing of non-pathological excisional injury to normal skin. However, based on the effects on fibroblast motility (e.g. Figs. 7 & 8), future work might explore whether the peptide also increases keratinocyte motility rate –thereby providing a contributory explanation of the accelerated re-epithelialization of pathologic skin wounds seen in response to αCT1 in humans.

The IAF hairless guinea pig^35^ demonstrated utility as a model in the present study. We moreover established approach based on wound splinting to generate a more human-like mechanical environment for our experiments on the scar reduction drug. To our knowledge, this is the first published report of the use of the IAF hairless guinea pig in an excisional skin injury or as a splinted wound model of scar formation. To date, this model has primarily been used in dermatologic studies of cutaneous drug, and allergen and/or toxin absorption.^71–73^ The guinea pigs proved more acquiescent to long-term experimentation, with most not disturbing their wound splints over the six-week duration of the study. The sparseness of hair follicles was also beneficial in maintaining wound splint adherence to the skin. In addition to favorable levels of docility and improved amenability to experimental manipulation, the IAF hairless guinea pig better replicated human αCT1 treatment results than the SD rat. Based on our favorable experience with this small animal, and the apparent relevance of results to human wound healing, splinted wounds on IAF hairless guinea pigs could be an excellent model for more efficient, clinically-relevant evaluation of the safety and efficacy of other wound healing therapeutics.

Limitations of the present work include the fact that the effects on αCT1 in humans and animal models studied herein were on small (<2 cm) wounds and in relatively mechanically quiescent regions of skin. Surgical procedures can involve large domains of injury, which may also be subject to dynamic, variable and heterogeneous patterns of strain during scar remodeling. It will be important to determine whether the effects of αCT1 on collagen organization and scar appearance are maintained in larger and more mechanically complex wound healing environments. At the cellular level, we did not directly link the changes in fibroblast migration induced by αCT1 to altered 3D patterns of collagen deposition. A tool that might assist with work of this type is GFP-tagged collagen and mouse lines expressing this protein.^74^ *In vitro* experiments are presently underway using live-cell imaging of fibroblasts derived from GFP-collagen expressing mice to observe patterns of collagen laid down by motile cells and determine whether αCT1 increases the disorganization of newly secreted ECM. A further limitation is that we do not understand the molecular bases of the effect of the peptide on dermal collagen order *in vivo*. We recently reported that a key aspect of Cx43 CT mimetic peptides in modulating the cardiac injury response relates to effects on the phospho-status of Cx43.^24^ In heart ischemia reperfusion models, it was shown that αCT1, and related peptides, bind to a domain on the endogenous Cx43 CT called H2, and in turn, promote phosphorylation of a serine residue at position 368 of the molecule. It remains to be established whether a similar mode-of-action operates in skin injury and if so, how αCT1-induced changes in Cx43 phospho-status might be linked altered fibroblast motility and/or long-term scar collagen organization. Given that improvements in the appearance of scars by 10–15% are considered clinically meaningful,^75^ further insight into how, at cellular and molecular levels, αCT1 prompts a 47% improvement in the long-term visual appearance of surgical wounds would be a useful endeavor.

## Methods

### Peptide sequences

α–connexin carboxyl-terminal (αCT1) peptide corresponds to a short sequence at the Cx43 CT linked to an antennapedia internalization sequence (RQPKIWFPNRRKPWKKRPRPDDLEI) as first described and characterized in Hunter et al.^22^ Peptide comprising only the antennapedia portion of the αCT1 peptide sequence (RQPKIWFPNRRKPWKK) or reversed inactive sequence (RQPKIWFPNRRKPWKKIELDDPRPR) were used as controls. All peptides were synthesized by American Peptide Co. Inc. (CA, USA) or Peptron Inc. (South Korea).

### Histological Samples from the Phase 1 Clinical Trial

The Phase I clinical trial on the effect of αCT1 on human dermal wound scars was performed in Switzerland on 49 healthy human volunteers in a randomized, double-blind Phase II study.^15^ The study protocol was approved by all institution ethics committees, was conducted in compliance with the principles of the Declaration of Helsinki and International Conference on Harmonization of Technical Requirements for Registration of Pharmaceuticals for Human Use (ICH) guidelines and Swiss regulations for clinical trials with medicinal products. Participants were notified of potential risks and benefits, were given the option to withdraw at any time, and signed informed consent forms before enrollment. On day 1 of the study a biopsy punch was used to create a circular 5 mm full-thickness (ie, stratum corneum to sub cutaneous fat layer) wound of unblemished skin underneath both arms (Fig. 1A). Wounds were internally randomized within each patient and one wound was treated with αCT1 in a hydroxyethylcellulose gel, while the other was treated with a vehicle control gel, enabling within-patient comparisons. Participating individuals (49 healthy subjects total) were randomized into four cohorts of 12-13 patients, wherein most of the subjects (41) their wounds were treated with vehicle gel or 20 μM (Cohort 1), 50 μM (Cohort 2), 100 μM (Cohort 3), or 200 μM (Cohort 4) of αCT1-containing gel. Ten patients were treated with matching vehicle control gel on both arms. Gel was applied immediately after injury and again 24 hours later. On day 29 of the study, a circular 2 mm full-width biopsy was sampled from the healed scar/granulation tissue formed at the two wound sites on each patient. One subject dropped out before completion of the trial. A total of 37 paired within-patient treatment and control skin samples were available for histological analyses. (Cohort 1 = 10, Cohort 2 =9, Cohort 3 = 8, Cohort 4 =10; n All Cohort_s_=37). The 100 μM dosage of αCT1 in Cohort 3 was the same dosage used in the Phase II clinical trial and considered the therapeutic dosage. The wounds were allowed to heal out to 29 days, with photographs taken of the healing wounds at regular intervals. At 29 days, the healed scars were photographed one last time and then biopsied. Biopsied scar tissue was washed, placed in 4% paraformaldehyde for 24 hours, and then embedded in paraffin for histology.

### Animal Wound Healing Models

All procedures were performed in accordance with the Guide for Use of Experimental Animals and IACUC committee at Virginia Tech (Virginia, USA) and conformed to the NIH Animal Care and Use Guidelines.^76^ Adult male Sprague-Dawley rats (n=15) were obtained from (Charles River Laboratories, USA). Four days before wound surgery, animals were anesthetized and dorsal fur removed by #7 size clippers followed by Nad’s (SI&D Inc., Garden Grove, CA USA) and VEET (Reckitt Benckiser LLC, USA), brand cold wax strips. The Nad’s strips were used first, to remove the majority of the fur, and then VEET strips (which are more adhesive) were used to remove the remaining strands. The waxed dorsal area was then treated with olive oil to remove excess wax and moisturize the skin to alleviate any irritation. The waxed area was gently covered with a piece of gauze and a commercial rat jacket (Lomir Biomedical Inc, USA), along with a custom-made wound covering was secured on the rat to prevent the animals interfering with their wounds. Adult male Institue of Armand Frappier (IAF) hairless guinea pigs weighing approximately 300 g (n=15) were obtained from Charles River Laboratories (Wilmington, MA USA.). Although mostly hairless, much of dorsum on these animals is covered with fine hairs. Two days prior to surgery, animals were anesthetized and hair was removed using Nad’s waxing strips, in order to create a cleared surface for surgically applying the splints. Olive oil was used to remove excess wax and relieve any skin irritation incurred. Animals were allowed to recover from anesthesia on a heating pad and then returned to group housing. In some cases, jackets (Lomir Biomedical Inc, USA), were also used for guinea pigs, though the majority of guinea pigs did not disturb their splints and thus did not require the jackets. On the day of surgery, animals were anesthetized with isoflurane (Henry Schein Animal Health, USA). placed on a prewarmed isothermal pad, and given 0.2 mg/kg (rats) or 0.03 mg/kg (guinea pigs) buprenorphine SR™-LAB (ZooPharm, USA) for analgesia. The dorsal region was prepped via 6 alternating scrubs of betadine (Dynarex Corporation, USA) and 70% alcohol. Six full thickness excisional wounds, 3 on either side of the spine, were created in the dorsal surface of each animal (Supplemental Fig. 3). 8 mm Miltex disposable biopsy punches (LifeSciences Inc., USA), were used to generate the wounds. Silicon splints (Bio-Labs Inc., USA) were cut using a Mayhew Pro 66000 hollow punch set (Mayhew Steel Products Inc., USA.). Sterile splints with inside diameter approximating the wound edge were secured with Krazy glue (Elmer’s Products Inc, USA) around the wound, and subsequently sutured through the full thickness of the skin with at least 6 interrupted 4-0 nylon sutures with a P-13 needle size (AD Surgical, USA). Replicating the methods used in cohort 3 of the Phase I clinical trial of αCT1, 100 μM of αCT1 was suspended in a 0.4% hydroxyethylcellulose gel (Natrosol, Ashland, USA) solution for the active treatment. Each animal received approximately 75 μL of gel per wound immediately after surgery and again 24 hours later; 3 wounds on one side of the spine received the active treatment, while the 3 wounds on the other side of the spine received control hydroxyethylcellulose gel with no peptide (Supplemental Fig. 3). The side of the spine that received treatment was randomized between animals. One paired set of active and control healed wounds were biopsied from each animal at 2-, 4-, and 6-weeks using a 10 mm biopsy punch. Animals received 4 mg/kg carprofen (Zoetis, USA) or 0.2 mg/kg buprenorphine for analgesia prior to the biopsies at 2 and 4 weeks. Animals were euthanized at 6 weeks. Biopsied scar tissues were washed, fixed in 4% paraformaldehyde for 24 hours, embedded in paraffin, sectioned across the midline of the scar at 10 μM and sections mounted on glass slides (2 sections/slide).

### Histochemistry and Histological Image Analysis

Biopsy sections were dewaxed and stained with Hemotoxylin and Eosin (H&E) or Picrosirius (PS) red using standard procedures. All assessments on either H&E stained or PS-red stained sections from either human and animal models were undertaken by investigators blinded to treatment or cohort. Measurements of epidermal thickness, dermal-epidermal junction length, epidermal length and counts of epidermal rete peg number, sub-epidermal melanocyte density, dermal nuclear density, dermal blood vessel density, sebaceous glands, hair follicles, and eccrine glands were carried out on images field-stitched (by Aperio Imagescope software, Leica, Germany) from images of human skin sections taken using a 20× objective on a Leica DM LB light microscope using ImageJ and approach similar to that we have previously described in Ghatnekar et al 2009.^29^ The scans included the full width (5-8 mm) of unwounded skin (e.g. Fig. 2A) and scar biopsies from both arms of all patients from the Phase I clinical trial, i.e., 2 wound locations x 2 time points x 37 patients = 148 scanned images of full-width skin/scar. For PS-red stained sections, collagen order was first assessed using automated scanning on an Olympus VS120 scanning microscope (Olympus Corporation, Tokyo, Japan) birefringence at a single polarization angle using standard methods. However, to capture the full breadth of the collagen fibers present in tissues sections, an Arduino-based microscope-addition capable of rotating the microscope condenser and analyzer in unison was constructed (Supplemental Fig. 2). A user interface system allowed the microscope user to specify a condenser angle, upon which point step motors actuated gears, 3D printed with a MakerBot Replicator 3D printer (MakerBot Industries LLC, USA), that interface with the microscope’s polarization condenser and rotatable analyzer. Slides were serially imaged at six polarization angles - 0°, 15°, 30°, 45°, 60°, & 75° - in order to obtain the full array of collagen fibers present in the sample with some overlap. A Matlab program^77^ was written to combine these six separate images and process them as one sample, using the MatFiber function^32^ to tally the fiber angles present in each image. Samples were analyzed along the depth of the tissue for collagen fiber density, local circular variance, local circular variance adjusted for local density (“collagen disorganization”), and standard deviation of collagen disorganization (“collagen disorganization variance”). Further details on the computational approach to measurement can be found in the Virginia Tech 2019 PhD thesis of Jade Montgomery.^78^

### *In Vitro* Analysis of Fibroblast Motility

#### Scratch Wound Assay

NIH-3T3 fibroblasts (ATCC, USA) were cultured in DMEM (Mediatech, USA) supplemented with FBS (Atlanta Biologicals, USA), 10U/ml penicillin and 10μg/ml streptomycin (Cambrex, USA), and 2mM L-alanyl-L-glutamine (Corning, USA) and kept in an incubator at 37°C, 5% CO_2_. For the assay, fibroblasts were plated on 6-well cell culture plates (TPP, USA) at 2×105 cells per well for 24 hours. Cells were treated with 1, 30, 60, 120, or 180μM αCT-1, vehicle (Veh; supplemented DMEM) or Antennapedia peptide for 24hrs at culture conditions. Cells were then scratched with a 200μl pipette tip, washed with PBS (Sigma, USA), and the same amounts of peptide were re-administered. Cells were allowed to recover in culture conditions for a further 24hrs, then washed with PBS, fixed for 10min in 2% paraformaldehyde (Fisher Scientific, USA) at room temperature and blocked with 1% BSA (Fisher Scientific, USA), 0.1% Triton-X-100 (Fisher Scientific, USA) in PBS. Cells were stained with Hoechst 33258 (Sigma, USA), mounted, and imaged on a Leica DM LB (Leica, Germany) fluorescent microscope equipped with a 20x/0.50 NA water immersion objective. The number of cells in each image was quantified by nuclear staining in NIH ImageJ (National Institutes of Health, USA). Live-imaging of scratch-wounded cells: NIH-3T3 fibroblasts were plated on poly-L-lysine (Sigma, USA) coated 35mm glass bottom culture plates (MatTek, USA) at 2×10^5^ cells per well for 24hrs. Cells were treated with either vehicle control solution or 180μM αCT-1 for 24 hours at culture conditions, and scratch-wounded as above. For the last 6hrs of the 24hr recovery period the media was replaced with CO_2_ independent media (Gibco, USA), and cells were imaged on an Axiovert 200M microscope (Carl Zeiss, Germany) equipped with a 10x/0.3 NA phase-contrast air objective (Carl Zeiss, Germany). Digital phase-contrast images were captured every 3 minutes by an Orca ER CCD camera (Hamamatsu) using Openlab 5.0.1 software (Improvision, USA).

#### In Vitro Cell Tracking Analysis

To test the potential effect of αCT on cell migration patterns, we tracked and analyzed the migration paths of NIH 3T3 fibroblasts plated on 2D tissue culture plastic as above for scratch wound assays. Live cell imaging was carried for eight hours under six different experimental conditions: no treatment control, 3 different dosages of 10, 50, and 100 μM αCT peptide, 100 μM reversed inactive control peptide, and 100 μM antennapedia inactive peptide controls. Using x-y coordinates at each time step, we calculated migration directions as the inverse tangent of y-displacement over x-displacement, then calculated angle changes as the difference between migration directions of subsequent time intervals. These angle changes ranged from −180° to 180° and represented how persistent cell migration followed along the same direction. To quantify the central tendency of cell directional persistence, we calculated the number of angle changes ≤30° as well as the mean vector length (MVL) of the angle change distribution according to Eq. 1:

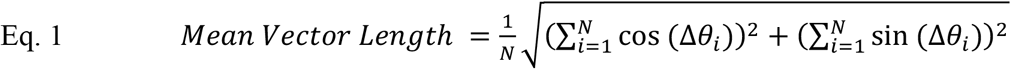

Where *Δθ_i_* is each angle change and *N* is the total number of angle changes. MVL quantified the alignment around a central angle and theoretically ranged from 0 (corresponding to no alignment) to 1 (corresponding to perfect alignment).

#### Computational Simulations of Wound Matrix Remodeling

To computationally analyze the potential effect of αCT1 on tissue-level collagen organization, we adapted a previously published agent-based model of fibroblast-collagen interactions.^41^ The full details of the computational approach have been discussed in prior studies; here we outline the model highlights. Briefly, we simulated a population of fibroblasts as individual agents that follow phenomenological rules capturing cell proliferation, apoptosis, orientation, migration, collagen deposition, and collagen degradation, all of which are functions of localized chemical, structural, mechanical, and persistence cues. The simulation geometry consisted of a 5mm deep x 5mm wide square wound surrounded by 3mm of non-wounded dermal tissue on the lateral and sub-dermal sides. The geometry was discretized into 10 μm x 10 μm collagen-containing elements, and collagen fiber structure (density and orientations) was tracked within each element as a distribution of fibers oriented in bins from −90 to 90 relative to the horizontal direction (parallel to the skin surface). Initially, the wound zone was devoid of all cells and matrix, while the surrounding non-wound zone was seeded with 9000 cells and collagen structure that matched reports of normal dermal tissue.^45^

Each cell was simulated as a disc of 5 μm radius that iterated through the following processes at each time step: (1) assess localized chemokine concentration and gradient vector, assess localized mechanical strain vector, assess localized collagen fiber orientation vector, assess cell persistence vector; (2) average the directional cues using a weighted vector resultant approach described previously,^38^ and reorient according to a circular probability distribution based on the average cue direction; (3) migrate in the newly reoriented direction at a prescribed migration speed; (4) deposit new collagen fibers parallel to the cell’s orientation at a prescribed deposition rate; (5) degrade a prescribed fraction of all fibers; (6) update the cell’s internal age clock and divide or die at appropriate intervals. These actions were repeated for each cell at each time step over a 28-day simulation period. Cell proliferation rate, cell migration speed, collagen deposition rate, and collagen degradation rate were all computed as functions of local chemokine concentration and set to match values reported in the literature.^37^

To simulate the effect of αCT on cell directional persistence, we multiplied the strength of the cell orientation probability distribution by an adjustment factor that allowed us to turn down a cell’s sensitivity to its local directional cues. This adjustment made each cell’s orientation decision slightly more variable with the αCT1 treatment vs. untreated (i.e., Vehicle) control simulations. Based on the effect of αCT1 on *in vitro* cell migration tracks in this study, we turned this scaling factor to 0.32 for the first 14 days of the healing period then ramped the factor back up to normal for the remaining 14 days in order to capture the dynamic drug treatment time course likely to occur *in vivo*.

### Statistical Methods

For statistical analysis of histological data, all depth-wise variables extracted from the whole-section images were analyzed in JMP^®^ Pro 12. To control for within-patient variance and any potential depthwise effects, a mixed-model analysis was used. Patient/subject designator was set as a random effect, while treatment, depth and their interaction were set as fixed effects. Due to the parametric requirements of mixed-model analysis, data was minimally transformed when necessary to achieve a homoscedastic normal distribution of residuals. Individual treatment to control comparisons at each depth were conducted via Tukey HSD and Student’s t test. For the scratch-wounding migration assay, averages were determined from individual measurements from 5 separate experiments. Measurements were taken from 10 fields for each treatment group. For multiple comparisons ANOVA with post-hoc analysis was performed. All statistics were prepared in PASW Statistics version 17.0.2. For assays of fibroblast pivoting, Non-parametric Kruskal-Wallis and Dunn’s multiple comparison tests were used to compare αCT1 treatment groups with controls. For analysis of fibroblast mean vector lengths (MVLs), data was cosine and sin transformed, then summed and mean square was calculated. Deviations were calculated via the same transformations to ensure uniformity. Statistical significance was determined on transformed mean square values via a Kruskal-Wallis single comparison test or a Mann-Whitney U-test. For all tests p<0.05 was considered significant.

## Author Contributions

Conception and design: Robert G. Gourdie, Jade Montgomery

Development of methodology: Jade Montgomery, William J. Richardson, J. Matthew Rhett, L. Jane Jourdan, Jeffrey W. Holmes

Acquisition of data (provided reagents, provided facilities, etc): Jade Montgomery, William J. Richardson, J. Matthew Rhett, Francis Bustos, Christina L. Grek, Gautam S., Ghatnekar, Katherine Degen, L. Jane Jourdan

Analysis and interpretation of data (e.g., statistical analysis, computational analysis): Jade Montgomery, William J. Richardson, Spencer Marsh, Robert G. Gourdie

Writing of the manuscript: Robert G. Gourdie

Review and revision of the manuscript: All authors

## Acknowledgments

We thank Linda Collins for her editorial contribution to this article.

## Supplemental Figures

**Supplemental Figure 1.**
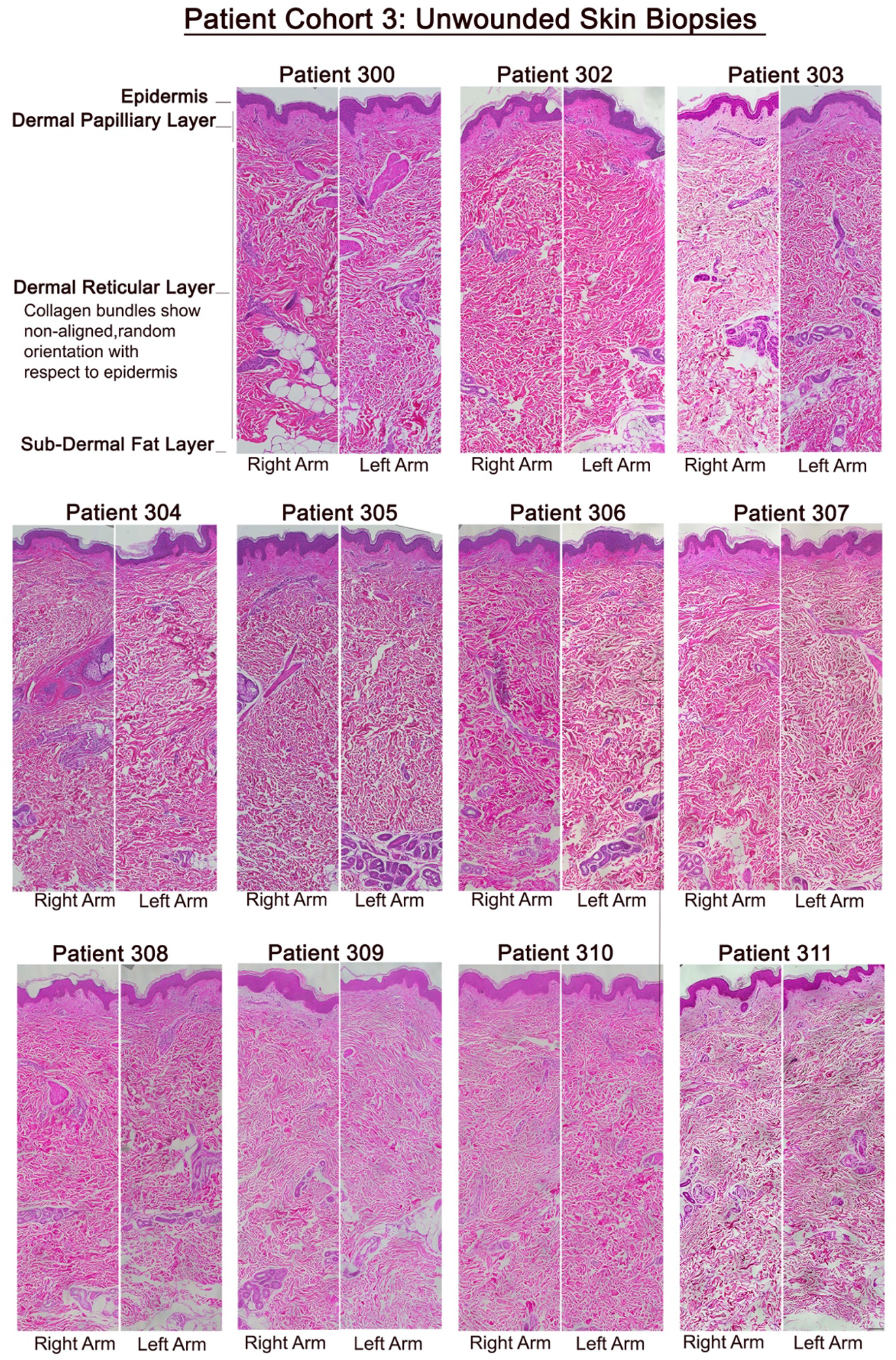
Unwounded skin sampled on day 1 of the Phase I clinical trial from the right and left arms of patients in cohort 3. The patterns were similar in unwounded skin in all patients sampled for the clinical trial, wherein dermal collagen bundles appear randomly aligned and arranged in sworls. Scale bar = 100 μm.

**Supplemental Figure 2.**
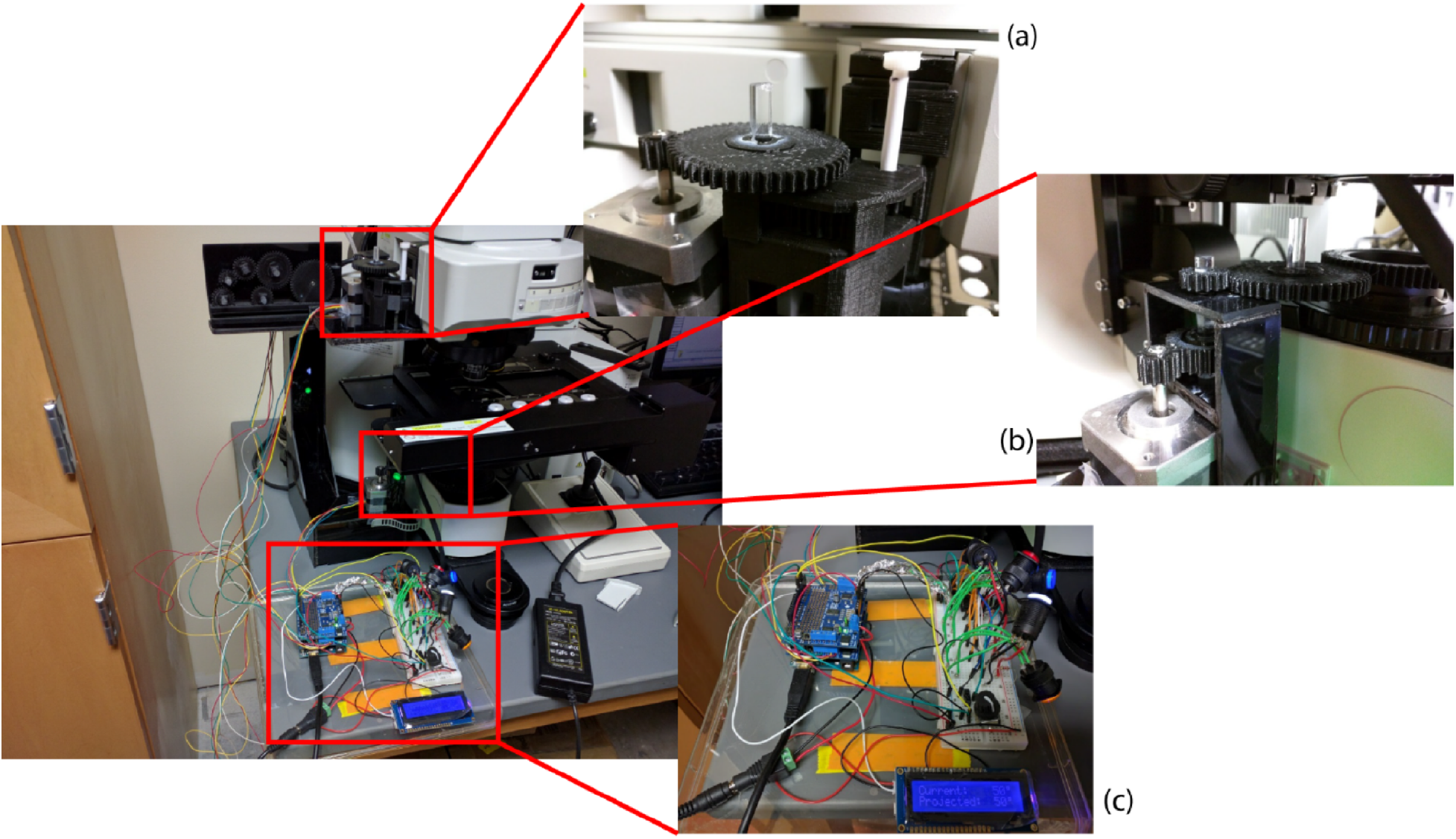
In order to capture the full breadth of the collagen fibers present in the system, we built and installed an Arduino-based^136,137^ microscope addition to our Olympus VS120 automated slide scanning scope capable of rotating the microscope condenser and analyzer in unison. The addition on the microscope allowed imaging at multiple consistent polarization angles. The left hand image shows the full addition in place on the Olympus VS120, with inset (a) showing the upper motor-analyzer interface, inset (b) showing the lower motor-condenser interface, and inset (c) showing the user control interface and Arduino controller system. A user interface system allows the microscope user to specify a condenser angle, upon which point step motors actuate gears 3D printed with a MakerBot Replicator 3D printer^138^ that interface with the native Olympus VS120 scanning microscope^122^ polarization condenser^139^ and rotatable analyzer^140^. Slides were serially imaged at six polarization angles - 0°, 15°, 30°, 45°, 60°, & 75° - in order to obtain the full array of collagen fibers present in the sample with some overlap. A MATLAB^123^ program was written to combine these six separate images and process them as one sample, using the MatFiber function^141^ to tally the fiber angles present in each image.

**Supplemental Figure 3.**
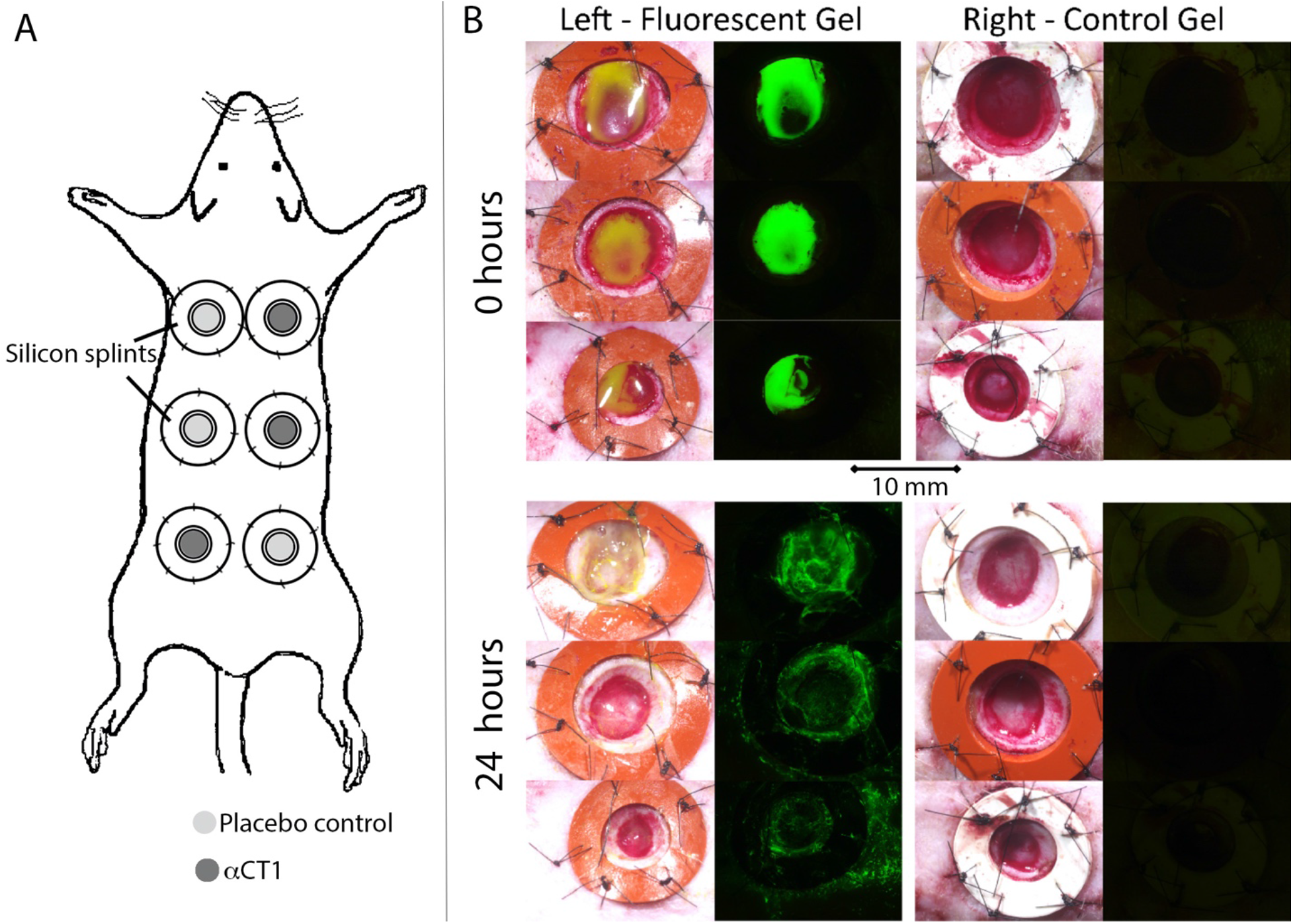
**A)** Experimental design used in the animal studies. The animal subjects (8 rats and 13 Guinea pigs) received six 10 mm circular wounds on their dorsal skin enabling within-subject comparisons of the effects of acute treatment with the therapeutic 100 μM dose of αCT1 relative to a vehicle control gel. Treatments were balanced within subjects and treated in a randomized, blinded manner. **B)** To maintain mechanical tension during healing, 8 mm thick circular silicon splints were placed around wounds to place uniform tension over the 6-week course of the study. As there were multiple wounds per subject we undertook a study of the diffusion of solutions from individual wounds. A fluorescent gel was placed in each wound directly after skin biopsy. 24 hours later fluorescence was only observed in wounds it was placed into, and not in wounds directly adjacent to the labeled wound.

## Supplemental Movies

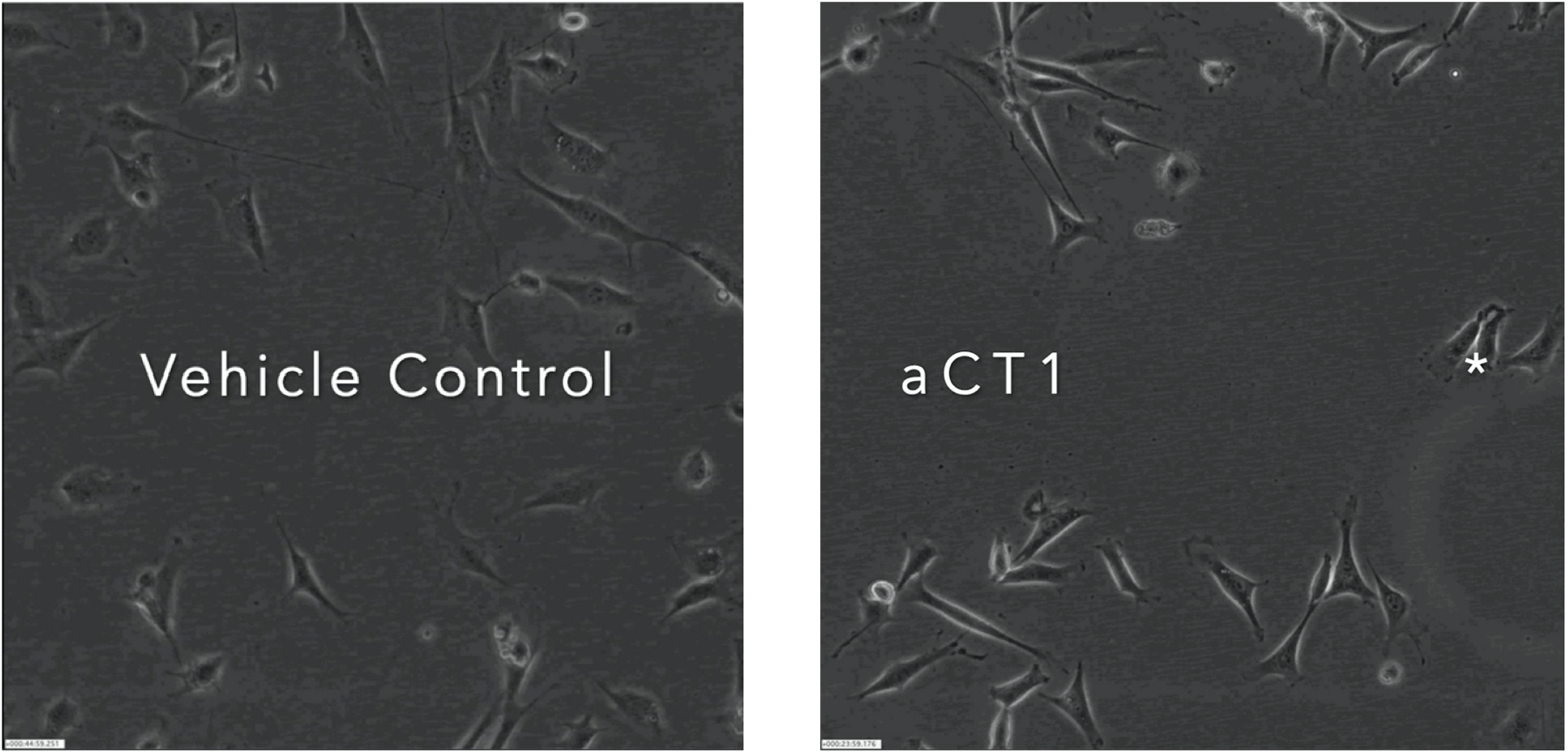
Video sequences of fibroblasts in a scratch wound migration assay 24 hours after scratching under vehicle control and 180 μM αCT1 treatment conditions. The sequences were generated on a Perkin Elmer live cell imaging microscope using phase contrast (10 x objective) optics. The microscope fields shown were taken at 5 minute intervals over 6 hours. A sequence of frames in Figure 7B shows the subsequent motions of the asterisked cell in the first few frames of the αCT1 sequence. This fibroblast exhibits a characteristic of αCT1-treated cells, wherein they demonstrate decreased directional persistence that is caused by repetition of a motility behavior involving cellular tail extension, release and pivoting.

